# An equine Endothelin 3 cis-regulatory variant links blood pressure modulation to elite racing performance

**DOI:** 10.1101/2022.11.04.515141

**Authors:** Kim Fegraeus, Maria K Rosengren, Rakan Naboulsi, Ludovic Orlando, Magnus Åbrink, Annika Thorsell, Ahmad Jouni, Brandon D Velie, Amanda Raine, Beate Egner, C Mikael Mattsson, Göran Andersson, Jennifer R.S Meadows, Gabriella Lindgren

## Abstract

A previous selective sweep analysis of horse racing performance revealed a 19.6 kb candidate region approximately 50 kb downstream of the Endothelin 3 (*EDN3*) gene. EDN3 and other endothelin family members are associated with blood pressure regulation in humans and other species, but similar association studies in horses are lacking. We hypothesized that the sweep region includes a regulatory element acting on *EDN3* transcription, ultimately affecting blood pressure regulation and athletic performance in horses. Selective sweep fine- mapping identified a 5.5 kb haplotype of 14 SNPs shared within Coldblooded trotters (CBT) and Standardbreds (SB). Most SNPs overlapped potential transcription factor binding sites, and haplotype analysis showed significant association with all tested performance traits in CBTs and earnings in SBs. From those, two haplotypes were defined: an elite performing haplotype (EPH) and a sub-elite performing haplotype (SPH). While the majority of SNPs in the haplotype were part of the standing variation already found in pre-domestication horses, there has been an increase in the frequencies of the alternative alleles during the whole history of horse domestication. Horses homozygous for EPH had significantly higher plasma levels of EDN3, lower levels of EDN1, and lower exercise-related blood pressure compared to SPH homozygous horses. Additionally, a global proteomic analysis of plasma from EPH or SPH homozygous horses revealed higher levels of proteins involved in pathways related to immune response and complement activation in the SPH horses. This is the first study to demonstrate an association between the *EDN3* gene, blood pressure regulation, and athletic performance in horses. The results advance our understanding of the molecular genetics of athletic performance, exercise-related blood pressure regulation, and biological processes activated by intense exercise.

**Author summary:** The horse is one of the most common species used for studying athletic performance. For centuries, horses have been used by humans for transportation, agriculture and entertainment and this has resulted in selection for various traits related to athletic performance. A previous study discovered that a genetic region close to the *Endothelin3* gene was associated with harness racing performance. Endothelin3 is known to be involved in blood pressure regulation and therefore we hypothesized that this region influences blood pressure and racing performance in horses. In this study we have used additional horses and fine-mapped the candidate region and we also measured blood pressure in Coldblooded trotters during exercise. Horses with two copies of the elite-performing haplotype had higher levels of Endothelin3 in plasma, lower blood pressure and better racing performance results, compared to horses with two copies of the sub-elite performing haplotype. We also discovered that horses with the sub-elite performing haplotype had higher levels of proteins related to the immune system in plasma. This study is the first to link Endothelin3 to blood pressure regulation and performance in horses. It broadens the understanding of the biological mechanisms behind blood pressure regulation as well as inflammation and coagulation system in relation to racing performance.

## Background

The use of domestic animals as models for genomic research has provided not only basic knowledge concerning gene function and biological mechanisms but also a complementary view on genotype–phenotype relationships (1). The horse is one of the most popular species used to study athletic performance. They have been intensively selected for centuries, to engender the optimal physical capacity for strength, speed, and endurance (2, 3). Additionally, their recent population history, involving closed populations with similar phenotypic traits within breeds and large variations across breeds, has created a favorable genome structure for genetic mapping. These factors combined make the horse an optimal model for studying the molecular genetics underlying athletic performance and the complex biological processes activated by exercise (1,2,4,5). Previous work in horses has started to reveal the effect of genes linked to complex traits such as muscle mass and locomotion patterns on athletic performance, and many potential candidate genetic variants have been identified (6–12). However, understanding the mechanisms by which the identified variants exert their functions is much more challenging.

A selective sweep study on harness racing breeds revealed a 19.6 kb genomic region on chromosome 22, located about 50 kb downstream of the *Endothelin 3* (*EDN3*) gene, under selection in Coldblooded trotters (CBTs) (13). Five SNPs in high linkage disequilibrium (LD r^2^ = 0.92–0.94) were significantly associated with racing performance, including the number of victories, earnings, and racing times. SNP NC_009165.3:g.46717860C>T (EquCab3.0) showed the strongest association within the sweep and was further genotyped in 18 additional horse breeds. The favorable T allele was found in high frequency in breeds used for racing, while it generally remained at low frequency in ponies and draught horses. It was hypothesized that the identified region might contain a regulatory element influencing either the expression of *EDN3* or other genes nearby (13).

The endothelin system encompasses the EDN ligands (e.g., EDN-1 to -3) and their G protein- coupled receptors (endothelin receptor type A and B). Together, they are involved in a broad range of biological functions, including cell proliferation and vasoconstriction (14). EDN2 and -3 can be considered as natural antagonists for EDN1 (15). The endothelin receptor type A (EDNRA) has the highest affinity for EDN1 and EDN2 but a low affinity for EDN3 (16). The endothelin receptor type B (EDNRB) has equal affinity for all three isoforms of endothelin (17). During exercise, endothelins are involved in the dynamic redistribution of blood flow to the lung and active muscles, limiting the flow to non-exercise-essential tissues, such as kidneys and intestines (15). Generally, EDNRA activation causes vasoconstriction and increased blood pressure, while EDNRB activation results in nitric oxide release, vasodilatation, and decreased blood pressure (18).

To date, most studies of the EDN system have focused on the role of EDN1. In human health studies, *EDN1* genetic variants were found to be implicated in hypertension (19). In performance trials, increased EDN1 plasma levels were observed in relation to exercise duration (20), with elevated blood pressure leading to impaired exercise capacity in elite athletes (20, 21). There have also been performance studies in horses, where EDN1 plasma levels largely mirrored the results found in humans, showing that the EDN1 response to exercise challenge is conserved across both species (22). There is a clear role for vasoconstriction in exercise performance, but the action of the EDN3 ligand is unclear. In this study, we aimed to fill this gap of knowledge. First through the fine-mapping of the *EDN3* region flanking the selective sweep, then the characterization of the spatial and temporal distribution of key genetic variants within horse populations, and finally by evaluating the effect of opposite haplotypes on blood pressure, as well as EDN1 and EDN3 plasma levels, before and during exercise. We also performed a global proteomic study on plasma to identify differentially expressed proteins amongst the various *EDN3* haplotypes. While the key breed of the study was the CBT, a popular harness racing breed in Sweden and Norway, the results are placed in the context of additional racing breeds. The results presented here on the endothelin pathway not only give new insights into genetic regulation of its established vasomodulatory properties, but also its emerging role in inflammation.

## Results

### Fine-mapping and comparative analyses suggest a regulatory role of the selected region

A 2018 study using the sparse Illumina SNP50 Genotyping BeadChip, identified a 19.6 kb selective sweep shared between SBs and CBTs (ECA22:46,702,297-46,721,892; Figure 1A). The sweep was located between *EDN3* and ENSECAG00000039543 (lncRNA), and five SNPs within this region were found to be associated with harness racing performance (13). From those five SNPs, g.46717860 C>T was the most significant. With a four-step process, we reduced this 19.6 kb sweep region to a minimal 5.5 kb region (Figure 1).

**Figure 1.**
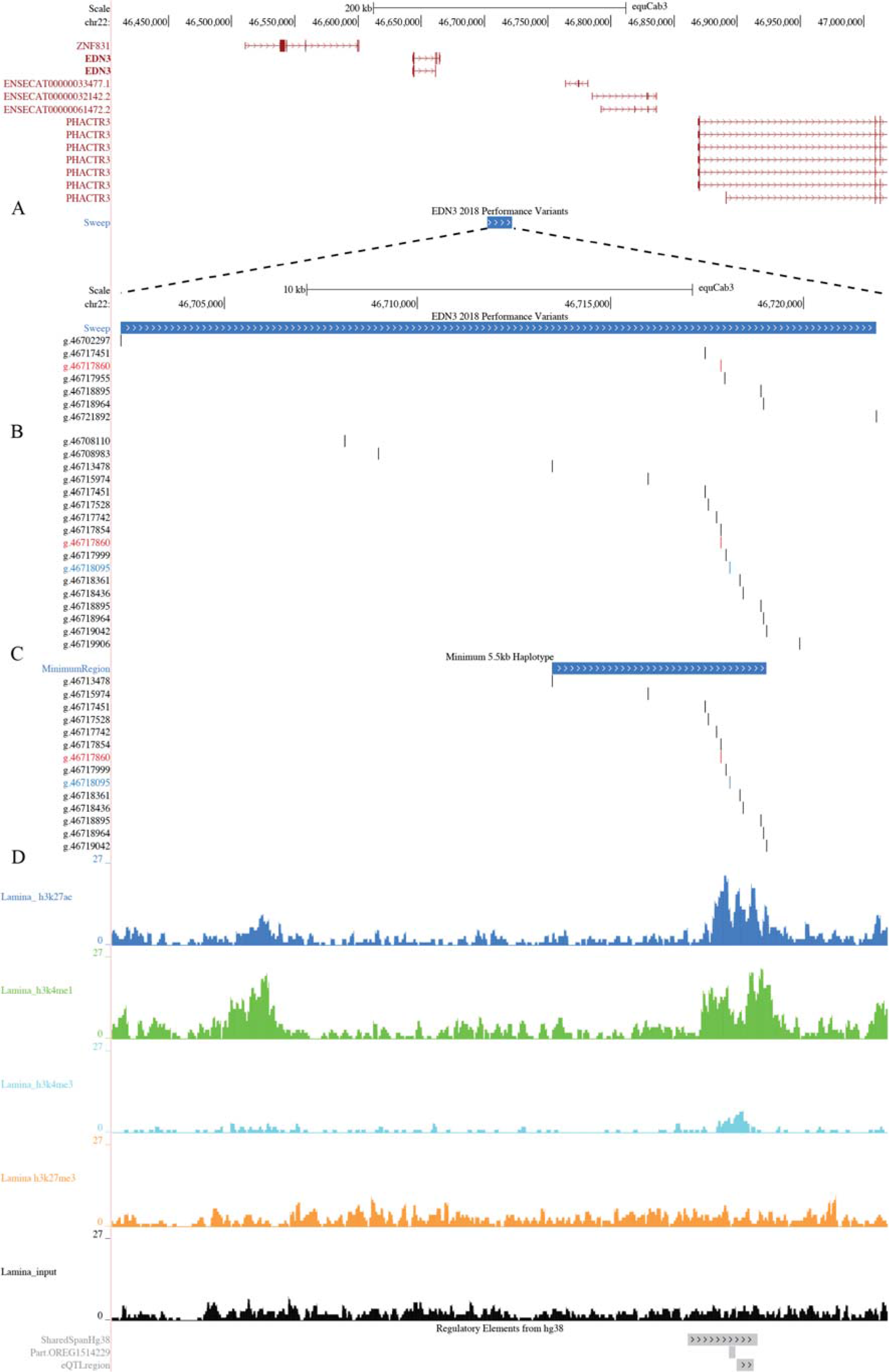
A. Overview of the 19.6 kb 2018 selective sweep region including all annotated genes in the region (13). B. Zoomed in view of the 2018 selective sweep region including the 17 SNPs used in the haplotype analysis. C. Overview of the 5.5 kb minimum shared haplotype within the Coldblooded trotters and Standardbreds, including all SNPs from the haplotype analysis. D. Horse functional data as generated by chip-seq of histone modifications from hoof lamina (from Equine Functional Annotation of Animal Genomes, FAANG) indicating an active enhancer in our region of interest. In addition, ∼ 2 kb of the associated haplotype region aligned with the human reference, including segments of an ORegAnno regulatory element, pinpointing a curated regulatory annotation of this region, (OREG1514229), GTEx cis-eQTL variants regulating *EDN3* in esophagus mucosa tissue.

In step one, we performed a sweep-performance association analysis using additional horse material (n=661) and SNP genotypes extracted from the 670K Axiom Equine Genotyping Array (Figure 1A). However, supplementing the 400 CBTs used originally in the 2018 study(13) with 221 additional CBTs did not provide increased resolution into the 19.6 kb sweep. Seven SNPs within the original sweep span were extracted from the array, including g.46717860. However, pairwise LD (r^2^ > 0.6) to g.46717860 resulted in the same five SNP set as found in (13), with all SNPs remaining significantly associated with harness racing performance traits in linear models (i.e., number of victories, earnings, and race times, P ≤ 0.05) (Supplementary Table S1).

**Table 1.**
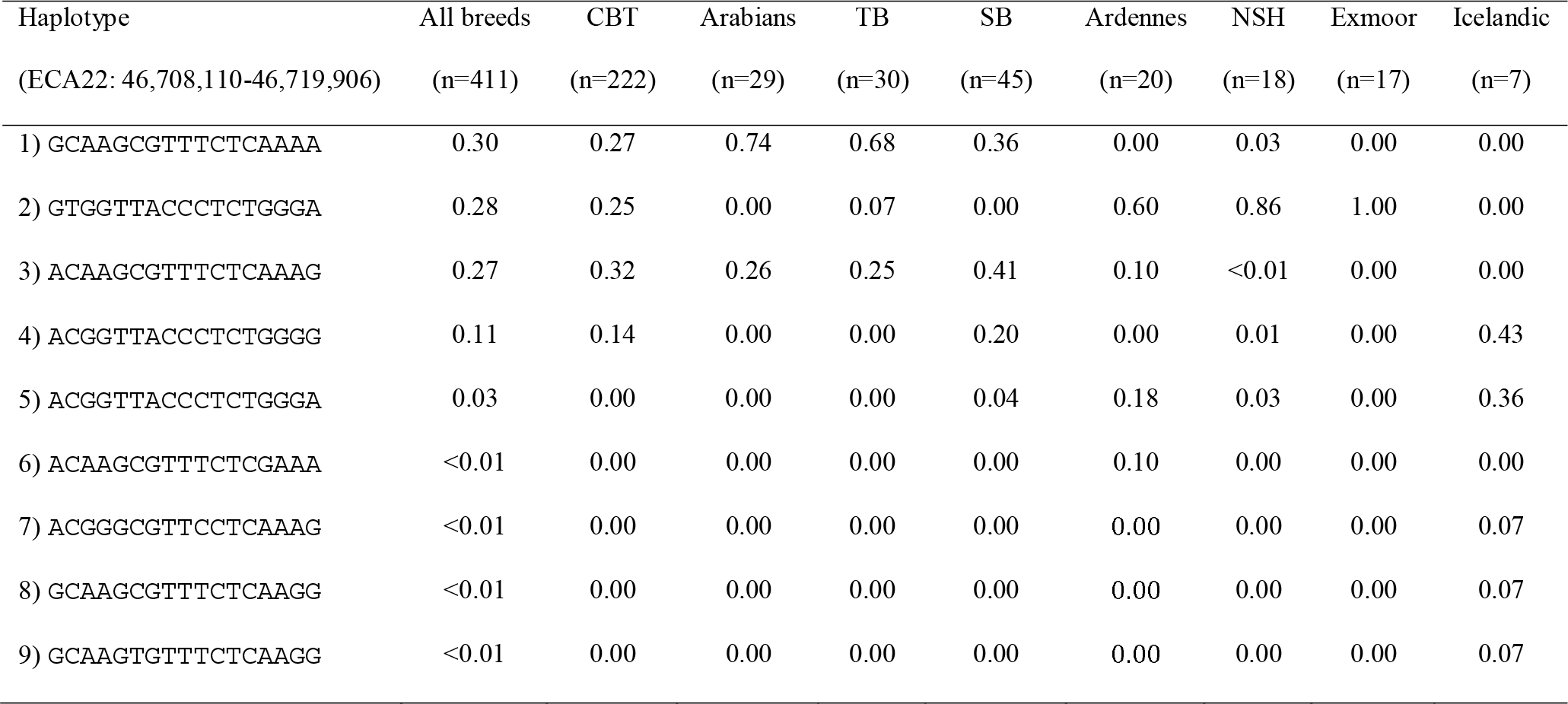
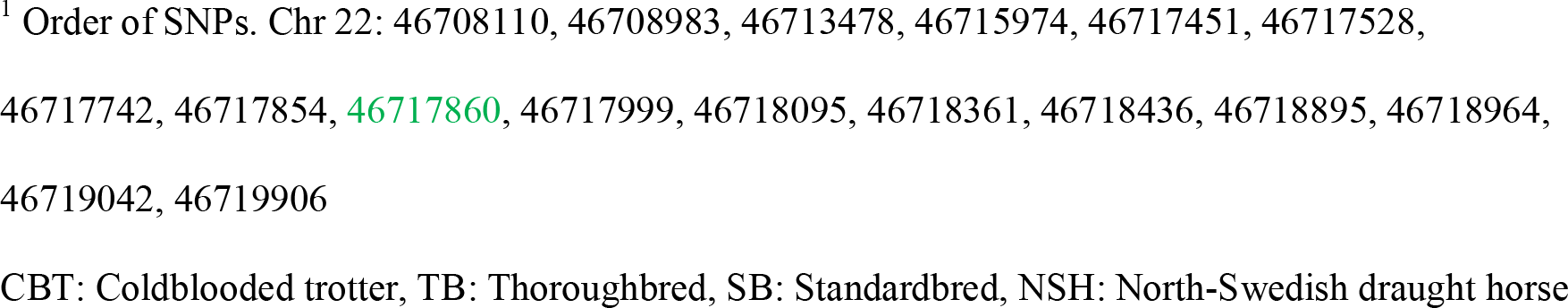
Haplotype frequencies in different breeds

In step two, we performed variant discovery to increase variant density across the 19.6 kb sweep region. Here, Illumina short-read whole genome sequences (WGSs) were generated from two CBTs and two SBs (the most common breed used for harness racing). In each breed, the horses were selected to be homozygous for different alleles at SNP g.46717860 (CC and TT, respectively) and to have either high (TT) or low (CC) earnings per start (Supplementary Table S2). From the 19.6 kb sweep region, 78 SNPs and six indels (one 400 bp deletion, one 23 bp deletion, three single bp deletions, and one single bp insertion) were identified. To possibly detect new structural variants, targeted long-read sequencing using Oxford Nanopore technology (ONT) was performed across the sweep region for eight horses: four CBTs with g.46717860-TT genotype and high performance, and four CBTs with the opposite features, including the two CBTs used for WGS (Supplementary Table S2). However, no new structural genetic variants were identified.

**Table 2.** Haplotype frequencies, haplotype coefficient and P-values for the performance analysis in Coldblooded trotters (n=180)

|  | Haplotype<sup>2</sup> (H3) |  | Haplotype (H1) |  | Haplotype (H2) |  | Haplotype (H4) |  |
| --- | --- | --- | --- | --- | --- | --- | --- | --- |
| Haplotype<sup>1</sup> | ACAAGCGT**T**TCTCAAAG |  | GCAAGCGT**T**TCTCAAAA |  | GTGGTTAC**C**CTCTGGGA |  | ACGGTTAC**C**CTCTGGGG |  |
| Frequency | 0.32 |  | 0.27 |  | 0.27 |  | 0.14 |  |
|  | Haplotype<br>coefficient | P-value<sup>3</sup> | Haplotype<br>coefficient | P-value<sup>3</sup> | Haplotype<br>coefficient | P-value<sup>3</sup> | Haplotype<br>coefficient | P-value<sup>3</sup> |
| No. of starts<sup>5</sup> | - | - | -0.30 | 0.43 | -1.00 | **0.004** | -1.05 | **0.02** |
| No. of wins<sup>5</sup> | - | - | -0.04 | 0.31 | -0.14 | **<0.001** | -0.17 | **<0.001** |
| No. of placings<sup>5</sup> | - | - | -0.06 | 0.13 | -0.12 | **<0.001** | -0.14 | **0.002** |
| Earnings (SEK)<sup>4,5</sup> | - | - | -0.31 | 0.05 | -0.53 | **<0.001** | -0.56 | **0.002** |
| Earnings/start<sup>5</sup> | - | - | -0.19 | 0.07 | -0.45 | **<0.001** | -0.42 | **<0.001** |
| Best time (sec/km)<sup>5</sup> | - | - | 0.03 | 0.16 | 0.07 | **<0.001** | 0.08 | **0.002** |

<sup>1</sup>Order of SNPs. Chr 22: 46708110, 46708983, 46713478, 46715974, 46717451, 46717528, 46717742, 46717854, **46717860**, 46717999, 46718095, 46718361, 46718436, 46718895, 46718964, 46719042, 46719906
<sup>2</sup>The most common haplotype was used as the base haplotype
<sup>3</sup>A GLM regression analysis was performed in R. Sex, age, birth country, number of starts (when applicable) and *DMRT3* genotype were included as fixed effects. Significant results ( $P \leq 0.05$ ) in bold
<sup>4</sup> SEK, Swedish kronor
<sup>5</sup> Log transformed values ( $\log_{10} + 1$ ) were used for wins and placings. Number of starts were square root transformed. Earnings and earnings per start were transformed applying $\ln(\text{Earnings} + 1000)$ .
CBT: Coldblooded trotter, TB: Thoroughbred, SB: Standardbred, NSH: North-Swedish draught horse

In step three, we prioritized variants for further analysis. While the whole genome sequenced elite CBT (high earnings per start) was homozygous for the 400 bp deletion (ECA22:46,714,602-46,715,003), the elite SB was heterozygous, and the deletion was not detected in the low-performing (low earnings per start) horses. The deletion was, thus, tested for association with racing performance traits in 479 CBTs using linear models, but the results were not significant (Supplementary Table S3). From the remaining variants identified, a total of 24 SNPs and one single base pair deletion, evenly distributed within the region, were selected (see methods) and genotyped in four donkeys and 412 horses sampled from 13 different breeds. SNP g.46717860 was not included in this set but was genotyped separately in 170 horses and imputed in the remainder. At this stage, 24 SNPs and 407 horses passed quality control measures. Pairwise LD estimation with SNP g.46717860 revealed 15 SNPs in strong LD (r^2^ ≥ 0.53, 14 SNPs with an r^2^ score > 0.98) (ECA22: 46,708,983- 46,719,042). LD decayed outside the region (r^2^ < 0.07), leaving 17 SNPs (15 SNPs in LD and two flanking SNPs) for haplotype analysis (Figure 1B). Phasing revealed 15 different haplotypes, five showing a frequency > 2 %. All haplotypes with a frequency > 5 % in the total sample set or within each breed are presented in Table 1 (disregarding breeds represented by less than five horses).

**Table 3.** Haplotype frequencies, haplotype coefficient and *P-*values for the performance analysis in Standardbreds (n=38)

|  | Haplotype<sup>2</sup> (H3) |  | Haplotype (H1) |  | Haplotype (H4) |  | Haplotype (H5) |  |
| --- | --- | --- | --- | --- | --- | --- | --- | --- |
| Haplotype<sup>1</sup> | ACAAGCGT**T**TCTCAAAG |  | GCAAGCGT**T**TCTCAAAA |  | ACGGTTAC**C**CTCTGGGG |  | ACGGTTAC**C**CTCTGGGA |  |
| Frequency | 0.43 |  | 0.34 |  | 0.18 |  | 0.04 |  |
|  | Haplotype | P-value<sup>3</sup> | Haplotype | P-value<sup>3</sup> | Haplotype | P-value<sup>3</sup> | Haplotype | P-value<sup>3</sup> |
|  | coefficient |  | coefficient |  | coefficient |  | coefficient |  |
| No. of starts<sup>5</sup> | - | - | 0.12 | 0.82 | 0.48 | 0.41 | -0.61 | **0.003** |
| No. of wins<sup>5</sup> | - | - | -0.02 | 0.74 | -0.08 | 0.35 | -0.47 | **<0.001** |
| No. of placings<sup>5</sup> | - | - | -0.03 | 0.63 | -0.11 | 0.06 | -0.25 | **<0.001** |
| Earnings (SEK)<sup>4, 5</sup> | - | - | -0.17 | 0.53 | -1.42 | **<0.001** | -1.32 | **<0.001** |
| Earnings/start<sup>5</sup> | - | - | -0.10 | 0.66 | -1.14 | **<0.001** | -1.12 | **<0.001** |
| Best time (sec/km)<sup>5</sup> | - | - | -0.10 | 0.37 | 0.15 | 0.24 | 0.65 | **0.01** |

<sup>1</sup>Order of SNPs. Chr 22: 46708110, 46708983, 46713478, 46715974, 46717451, 46717528, 46717742, 46717854, **46717860**, 46717999, 46718095, 46718361, 46718436, 46718895, 46718964, 46719042, 46719906 <sup>2</sup>The most common haplotype was used as the base haplotype <sup>3</sup>A GLM regression analysis was performed in R. Sex, age, birth country and number of starts (when applicable) were included as fixed effects. Significant results ( $P \leq 0.05$ ) in bold <sup>4</sup> SEK, Swedish kronor <sup>5</sup> Log transformed values (log10 +1) were used for wins and placings. Number of starts were square root transformed. Earnings and earnings per start were transformed applying ln(Earnings + 1000). CBT: Coldblooded trotter, TB: Thoroughbred, SB: Standardbred, NSH: North-Swedish draught horse

The association between haplotype and harness racing performance traits was performed in CBTs (n=180) and SBs (n=38) using generalized linear model regression (GLM) analysis in the statistical software R (23) (Table 2 and 3). Only haplotypes with a frequency > 2 % in the population were included in the analysis. Of the four haplotypes in CBTs, haplotype 3 (H3), carrying the g.46717860-T high performance associated allele, was the most common (0.32) and was defined as the base haplotype. Two haplotypes, H2 and H4, respectively, demonstrated a significant negative effect on all performance traits tested, and each carried the g.46717860-C low performance associated allele (Table 2). Four haplotypes were also available for testing within SBs with performance data. Three haplotypes were shared with CBTs (H1, H3 and H4) and one (H5) was shared with other breeds (Table 3). As for the CBTs, H3 was the most common haplotype (0.43) and was defined as the base haplotype in this analysis. Two haplotypes demonstrated significant negative effects on performance traits: H4 (earnings and earnings per start) and H5 (all traits tested). The above analysis revealed a minimum 5,564 bp, 14 SNP shared haplotype, significantly associated with racing performance within CBTs (ECA22: 46,713,478-46,719,042). The same variants were also significantly associated with racing performance in SBs. This allowed for the definition of an elite-performing haplotype (EPH: AAGCGTTTCTCAAA), and a sub-elite-performing haplotype (SPH: GGTTACCCTCTGGG) (Supplementary Table S4) (Figure 1C). Of note, the reference EquCab3.0 genome represents the SPH variant.

**Table 4.** Performance results in 539 Coldblooded trotters for SNP g.46718095 T>C

| Genotype | TT (n=65) |  | TC (n=256) |  | CC (n=218) |  | P<sup>2</sup> |  |  |
| --- | --- | --- | --- | --- | --- | --- | --- | --- | --- |
| Performance trait<sup>1</sup> | Mean | Median | Mean | Median | Mean | Median | TT/TC | TT/CC | TC/CC |
| No. of starts | 31.9 | 22.0 | 40.2 | 28.0 | 41.3 | 31.0 | **0.01** | **0.02** | 1.00 |
| No. of wins | 2.8 | 1.0 | 5.7 | 3.0 | 5.9 | 3.0 | **0.003** | **0.02** | 0.65 |
| No. of placings (1-3) | 9.3 | 4.0 | 14.2 | 9.0 | 14.5 | 9.0 | **0.005** | 0.07 | 0.38 |
| Earnings (SEK) | 207,295 | 51,772 | 438,700 | 140,161 | 453,911 | 115,312 | **0.006** | **0.05** | 0.58 |
| Earnings per start (SEK) | 4,752 | 2,976 | 9,666 | 4,362 | 8,469 | 3,771 | **0.003** | 0.08 | 0.20 |
| Best km time (sec)<sup>3</sup> | 90.9 | 91.4 | 88.5 | 88.3 | 88.9 | 88.9 | **0.001** | **0.02** | 0.44 |

<sup>1</sup>Log transformed values (log10 +1) were used for wins and placings. Earnings and earnings per start were transformed applying ln(earnings + 1000). Square root transformations were used for number of starts
<sup>2</sup>A linear model was performed in R. Sex, age, birth country, number of starts and *DMRT3* genotype were included as fixed effects (when significant). Least square means and Tukey were used as post hoc tests. Significant results ( $P \leq 0.05$ ) in bold
<sup>3</sup>Only races where the horse did not gallop were included

In step four, we assessed the potential functional impact of the minimal 5.5 kb region and each of the 14 SNP haplotype variants. Using horse functional data from two Thoroughbreds (from the Equine Functional Annotation of Animal Genomes, FAANG), we found three SNPs (g.46717860, g.46718095 and g.46718113) that overlapped potential active enhancers, i.e., H3K4me1 and H3K27ac marks (24, 25) (Figure 1D). Both Thoroughbreds were homozygous for EPH, with the histone marks indicating the presence of an active enhancer in the hoof lamina, present but less active in lung and ovarian tissue, and repressed in the skin(24, 25). In comparative analyses, a genome liftover to hg38 indicated that, although not well conserved, ∼ 2 kb of the associated haplotype region aligned with the human reference, including segments of an ORegAnno regulatory element (OREG1514229), GTEx cis-eQTL variants regulating *EDN3* in esophagus mucosa tissue (Figure 1D). These combined regulatory signals prompted us to test the potential for transcription factor binding affinity changes in relation to the EPH and SPH alleles. Most SNP allele changes (12 out of 14 SNPs) altered transcription factor binding (Supplementary Table S4). For EPH, this meant the perturbation of matrices for stress responders, e.g., IRF2 and NFKB1, while creating sites for developmental transcription factors such as NR4A2 and HOXA5.

### The elite performing haplotype has an additive effect on performance traits

Given that all 14 associated variants within the minimum shared 5.5 kb haplotype were in perfect LD, we used the genotypes at g.46718095 as a proxy for EPH (C allele) and SPH (T allele) haplotypic pairs and re-assessed the association between haplotypes and racing performance. For CBTs (n=539), EPH homo- or heterozygotes outperformed the SPH homozygotes (Table 4). In SBs, there were no statistically significant differences between the three genotypes (Table 5). However, when analyzing horses carrying at least one T allele versus CC horses, the results were significant for several performance traits (Table 5), including the number of wins, placings and earnings. However, the number of wins and placings was higher in horses with at least one T allele, while the CC horses earned more money than the TT/TC horses (Table 5).

**Table 5.** Performance results in 271 Standardbreds for SNP g.46718095 T>C

| Genotype | TT (n=7) |  | TC (n=62) |  | CC (n=202) |  | P<sup>2</sup> |  |  |  |
| --- | --- | --- | --- | --- | --- | --- | --- | --- | --- | --- |
| Perf. trait<sup>1</sup> | Mean | Median | Mean | Median | Mean | Median | TT/TC | TT/CC | TC/CC | CC/<br>TT+TC |
| No. of starts | 43.0 | 48.0 | 42.9 | 38.0 | 35.6 | 31.0 | 0.85 | 0.60 | 0.52 | 0.20 |
| No. of wins | 6.7 | 5.0 | 6.3 | 4.0 | 5.7 | 5.0 | 0.82 | 0.99 | 0.10 | **0.04** |
| No. of placings (1-3) | 15.6 | 12.0 | 16.1 | 16.0 | 13.7 | 11.0 | 0.86 | 0.96 | 0.08 | **0.03** |
| Earnings (SEK) | 700,008 | 322,300 | 730,124 | 332,300 | 1,075,529 | 369,675 | 0.92 | 0.43 | 0.06 | **0.01** |
| Earnings per start (SEK) | 11,981 | 5,463 | 16,673 | 9,765 | 35,346 | 9,897 | 0.86 | 0.44 | 0.16 | **0.04** |
| Best km time (sec)<sup>3</sup> | 73.1 | 73.9 | 73.3 | 72.8 | 73.1 | 72.9 | 0.85 | 1.00 | 0.19 | 0.10 |

<sup>1</sup>Log transformed values (log10 +1) were used for wins and placings. Earnings and earnings per start were transformed applying ln(earnings + 1000). Square root transformations were used for number of starts
<sup>2</sup>A linear model was performed in R. Sex, age, birth country and number of starts were included as fixed effects (when significant). Least square means and Tukey were used as post hoc tests. Significant results ( $P \leq 0.05$ ) in bold
<sup>3</sup>Only races where the horse did not gallop were included

Our racing performance analyses revealed that the g.46718095 genotypes did not distribute according to HWE. This prompted us to ask if there had been selection on this position or perhaps the minimal haplotype in other breeds. While each individual’s performance status is unknown, a trend for increased frequency of the C allele in traditional performance breeds was observed, e.g., Arabian horses, Thoroughbreds and Warmbloods. In contrast, this allele was low or absent in draft horses and ponies (Table 6).

**Table 6.** Allele frequency of SNP g.46718095 T>C

| **Breed** | **n** | **T** | **C** |
| --- | --- | --- | --- |
| Arabian horses | 30 | 0.00 | 1.00 |
| Thoroughbred | 30 | 0.07 | 0.93 |
| Standardbred | 306 | 0.14 | 0.86 |
| Warmblood | 3 | 0.17 | 0.83 |
| Coldblooded trotters | 539 | 0.36 | 0.64 |
| Finnhorses | 4 | 0.50 | 0.50 |
| Gotland pony | 2 | 0.50 | 0.50 |
| Shetland pony | 18 | 0.64 | 0.36 |
| Ardennes | 20 | 0.78 | 0.22 |
| Icelandic horses | 11 | 0.86 | 0.14 |
| North-Swedish draught | 25 | 0.88 | 0.12 |
| Exmoor pony | 23 | 1.00 | 0.00 |
| Fjord horse | 1 | 1.00 | 0.00 |
| Przewalski | 4 | 1.00 | 0.00 |

### Increased frequency of the favorable haplotype coincides with the time of horse domestication

For all 14 SNPs within the minimum shared haplotype in the trotters, we investigated the allele frequency over time using the mapDATAge package (26). For all SNPs but two (SNP 6 and 7 in Figure 2) there was an increase of the alternate allele from 7,500-5,500 years ago (Figure 2). For all SNPs considered, the two alleles segregated in specimens pre-dating the rise and spread of the DOM2 genetic lineage of modern domestic horses (data not shown). The dataset considered included 431 ancient genomes previously characterized (2,27–31).

**Figure 2.**
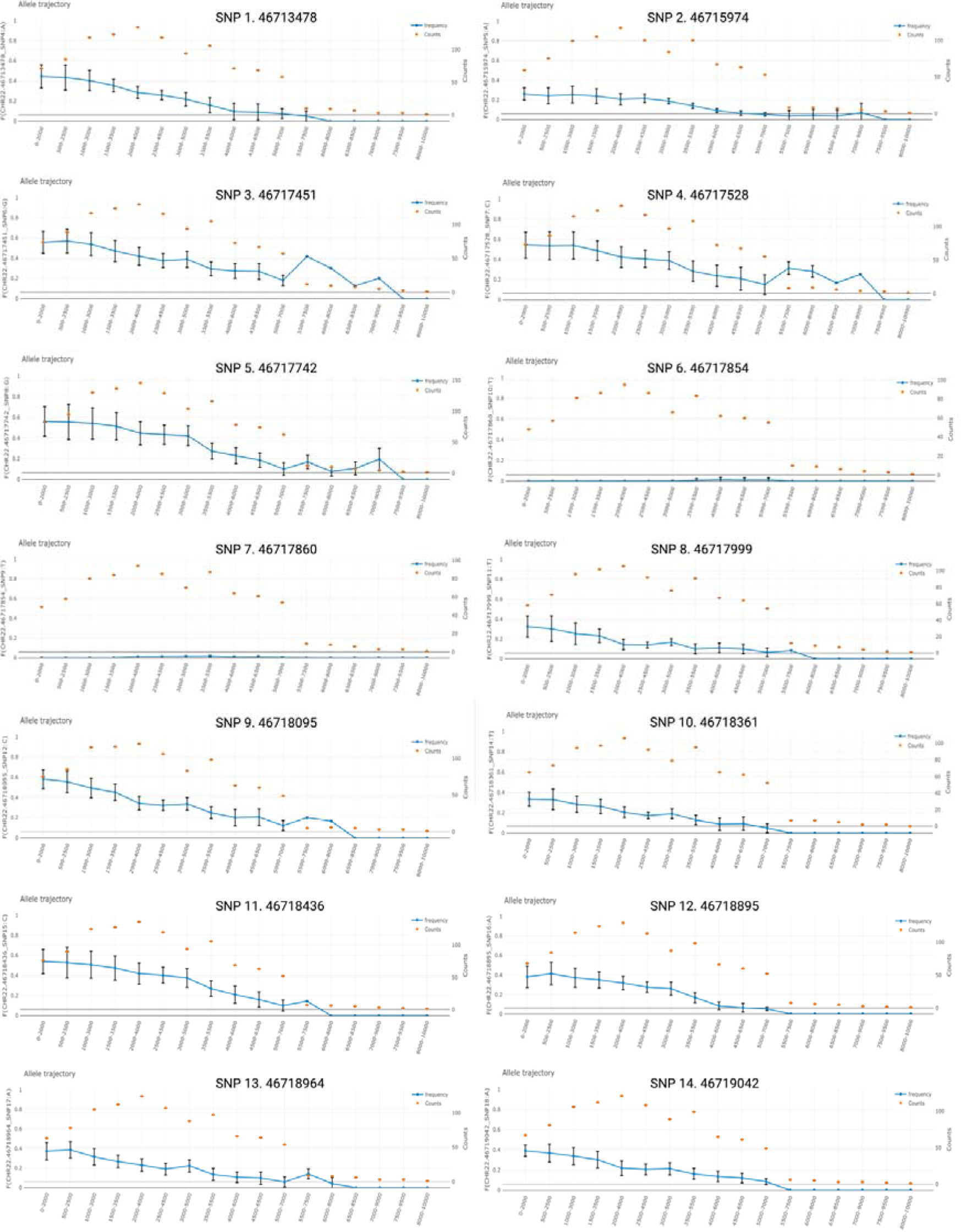
Frequency of the alternative allele over time from the mapDATAge package (26) for all 14 SNPs within the minimum shared haplotype in the trotters. The orange dots represent the number of horse samples considered in a given time bin of 2,000 years, and the blue line represents the frequency for the alternative allele estimated from allele counts. Figure created with Biorender.com.

### Elite performance haplotype linked to lower blood pressure

To link genotype to phenotype, we assessed the physiological difference between CBTs homozygous for SPH (n=5) and EPH (n=13) before, during and post-exercise (see methods). Blood pressure measurements were also taken from eight heterozygous horses (HET) at rest. Horses homozygous for SPH had, on average, significantly higher systolic blood pressure (SBP), diastolic blood pressure (DBP) and mean arterial pressure (MAP) during exercise, measured directly after the last uphill interval (Figure 3A-C). In addition, five mins after the uphill interval, the SPH group showed significantly higher SBP, MAP, and pulse pressure (PP) relative to the EPH group (Figure 3D-F). Horses homozygous for SPH also had significantly higher SBP and PP 20 mins after completing the exercise, compared to EPH horses (Figure 3G-H). Twenty mins after the exercise EPH horses were back to resting values, while it took ten additional mins for SPH horses to recover to resting values. There were no statistically significant differences in blood pressure measurements at rest, before the exercise. Furthermore, SPH horses had a higher heart rate than EPH horses during exercise (97.5 vs. 86.4) and five mins after exercise (85.7 vs. 77.0), although the difference was not significant. All blood pressure values are presented in Supplementary Table S5.

**Figure 3.**
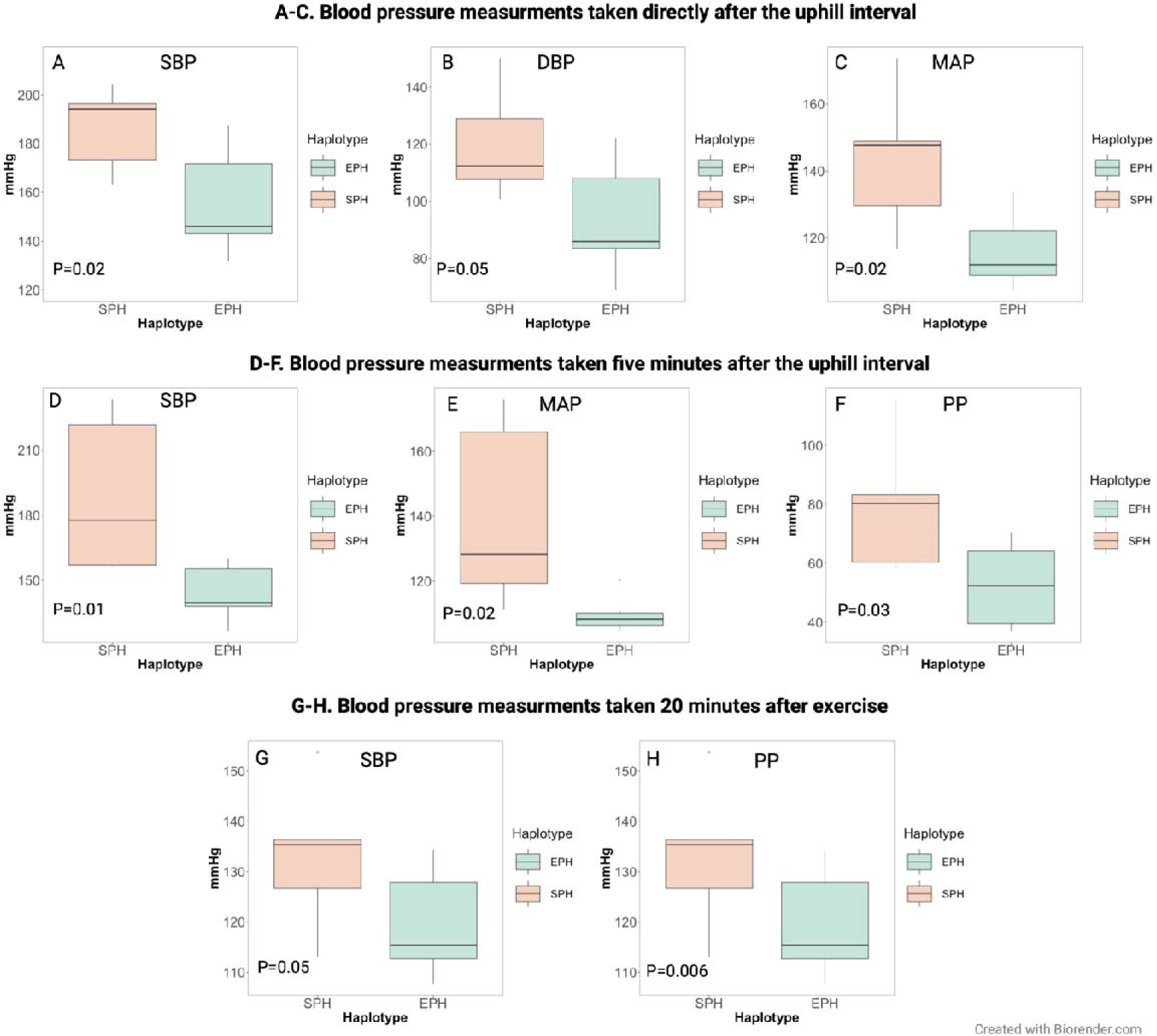
Blood pressure measurements during and after exercise in horses homozygous for the sub-elite performing haplotype (SPH) and horses homozygous for the elite-performing haplotype (EPH). A-C: Blood pressure during exercise measured directly after the uphill interval (SPH n = 5, EPH n=7). D-F: Blood pressure during exercise measured five mins after the uphill interval (SPH n = 5, EPH n=7). G-H. Blood pressure measured 20 mins after completing the exercise (SPH n = 5, EPH n=11). SBP: Systolic blood pressure, DBP: Diastolic blood pressure, MAP: Mean arterial blood pressure, PP: Pulse pressure. PP is defined as the systolic blood pressure minus the diastolic blood pressure.

### Plasma concentrations of EDN1 and EDN3 were significantly different between the haplotype groups

ELISA tests were used to measure the plasma concentration of EDN1 and EDN3 at rest and during exercise for each haplotype group (Figure 4). Horses homozygous for SPH (n=8) had a significantly higher plasma concentration of EDN1 both at rest and during exercise compared to the other groups (Tukeýs HSD test). In addition, SPH horses had a significantly lower plasma concentration of EDN3 at rest and during exercise (borderline significant) compared to horses homozygous for the EPH (Figure 4). Individual plasma values are presented in Supplementary Tables S6 and S7.

**Figure 4.**
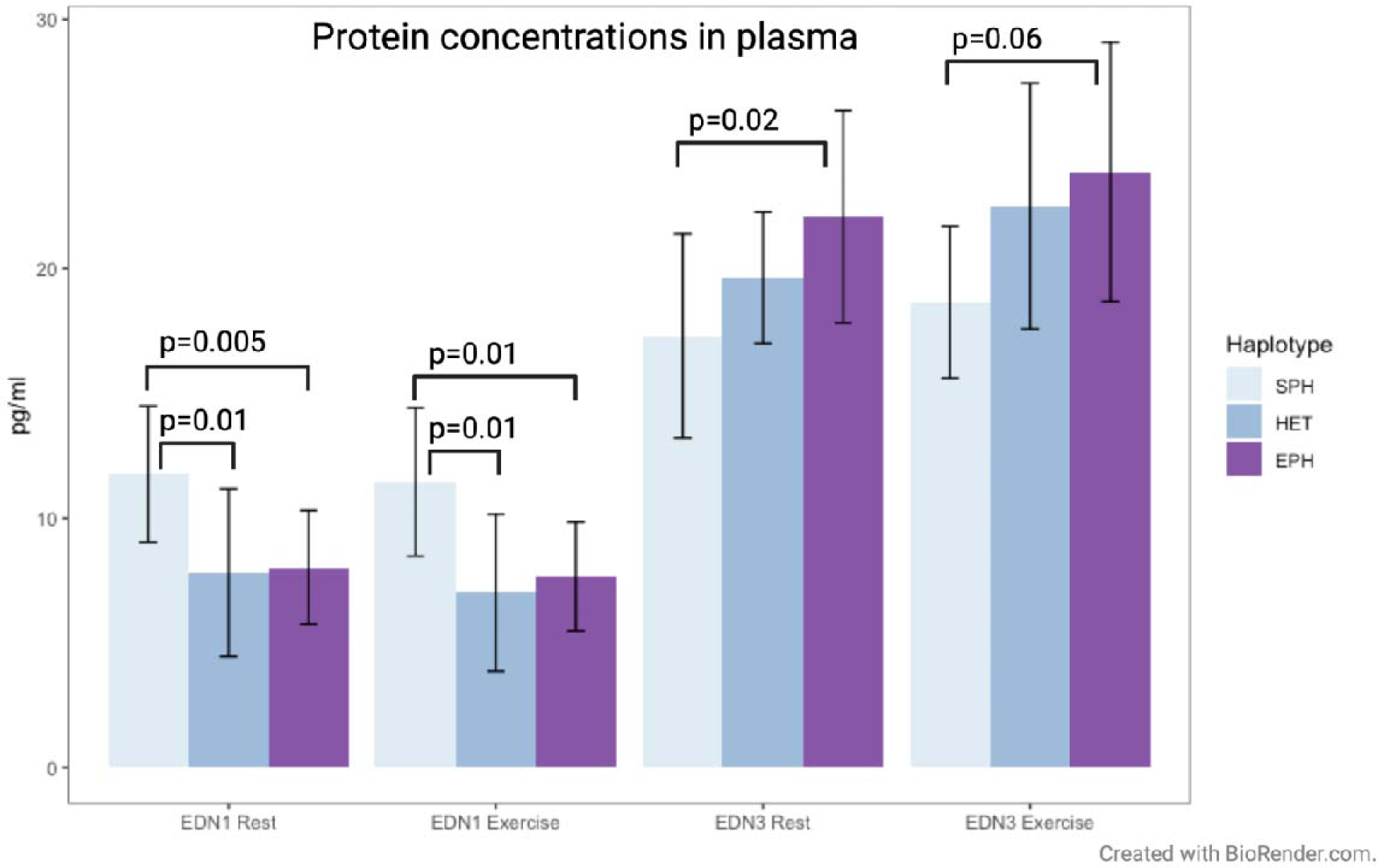
Plasma concentrations (pg/ml) of EDN1 and EDN3 at rest and during exercise in horses homozygous for the sub-elite performing haplotype (SPH) (n=8), horses heterozygous for the EPH and SPH (HET) (n=11) and horses homozygous for the elite-performing haplotype (EPH) (n=21). ANOVA and Tukeýs HSD test were performed. Standard deviations are presented as error bars.

### Significant differences in protein expression between haplotype groups

To identify potential biological pathways including EDN1 and EDN3, and to compare the expression of different proteins between EPH (n=6) and SPH (n=6) homozygous carriers, we performed a global relative quantitative proteomic analysis using Tandem Mass Tag (TMT) and mass spectrometry (MS), using plasma samples taken at rest or during exercise. In total, 582 distinct proteins were identified, but neither EDN1 nor EDN3 (Supplementary Table S8). Proteins quantified from at least three horses from each haplotype group were included in the statistical calculations. This resulted in 383 proteins that were analyzed in the five following comparisons; 1) exercise vs. rest in SPH, 2) exercise vs. rest in EPH, 3) EPH vs. SPH at rest, 4) EPH vs. SPH during exercise, 5) paired ratio analysis, where the individual difference at rest and exercise was compared between EPH and SPH groups.

Fifty proteins were found to show significantly different expression levels across the five comparisons (P < 0.05; Supplementary Table S8). Table 7 presents the top proteins with the highest fold change in each comparison, including all proteins with different levels that were significant in more than one comparison.

**Table 7.**
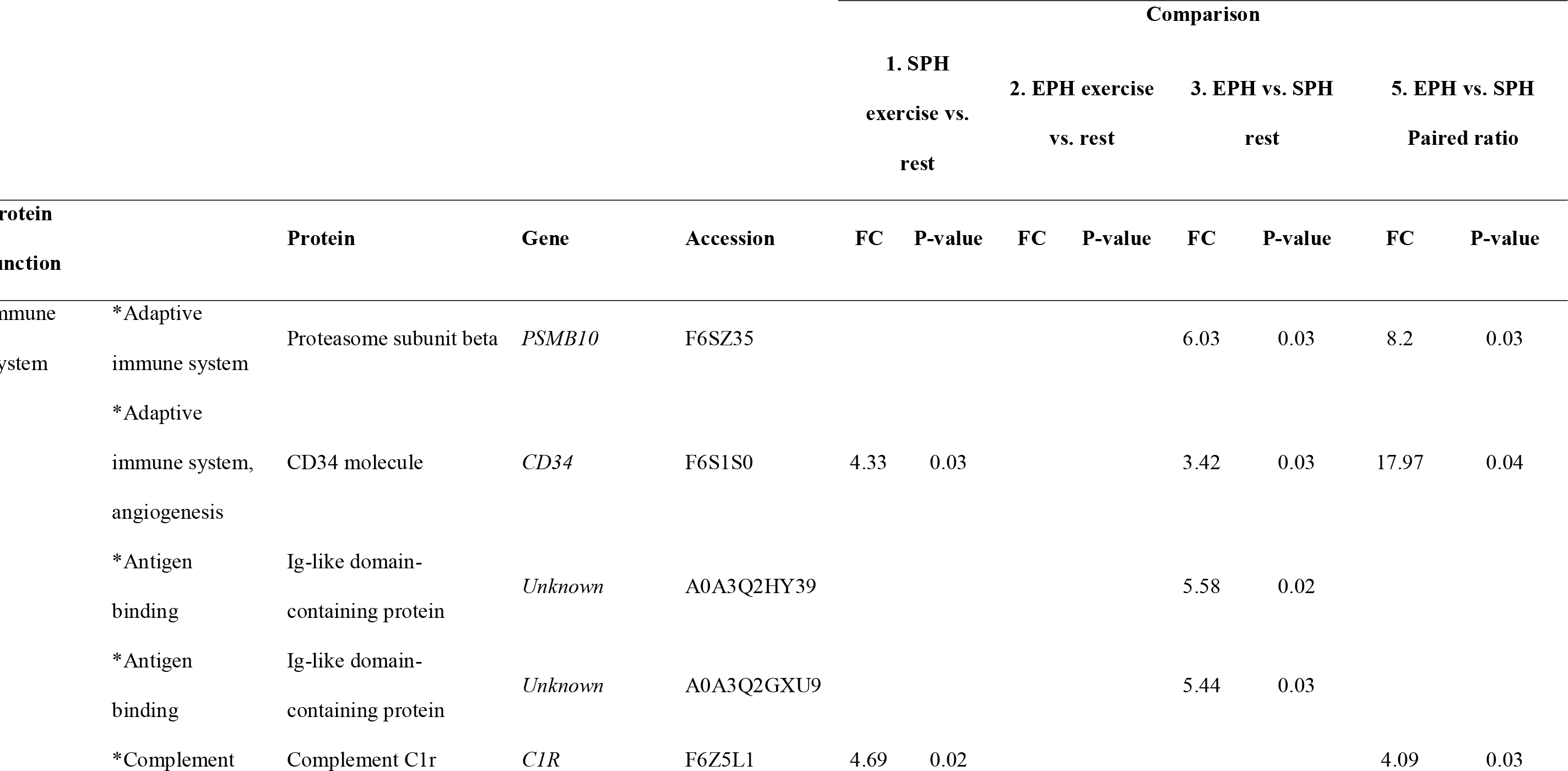
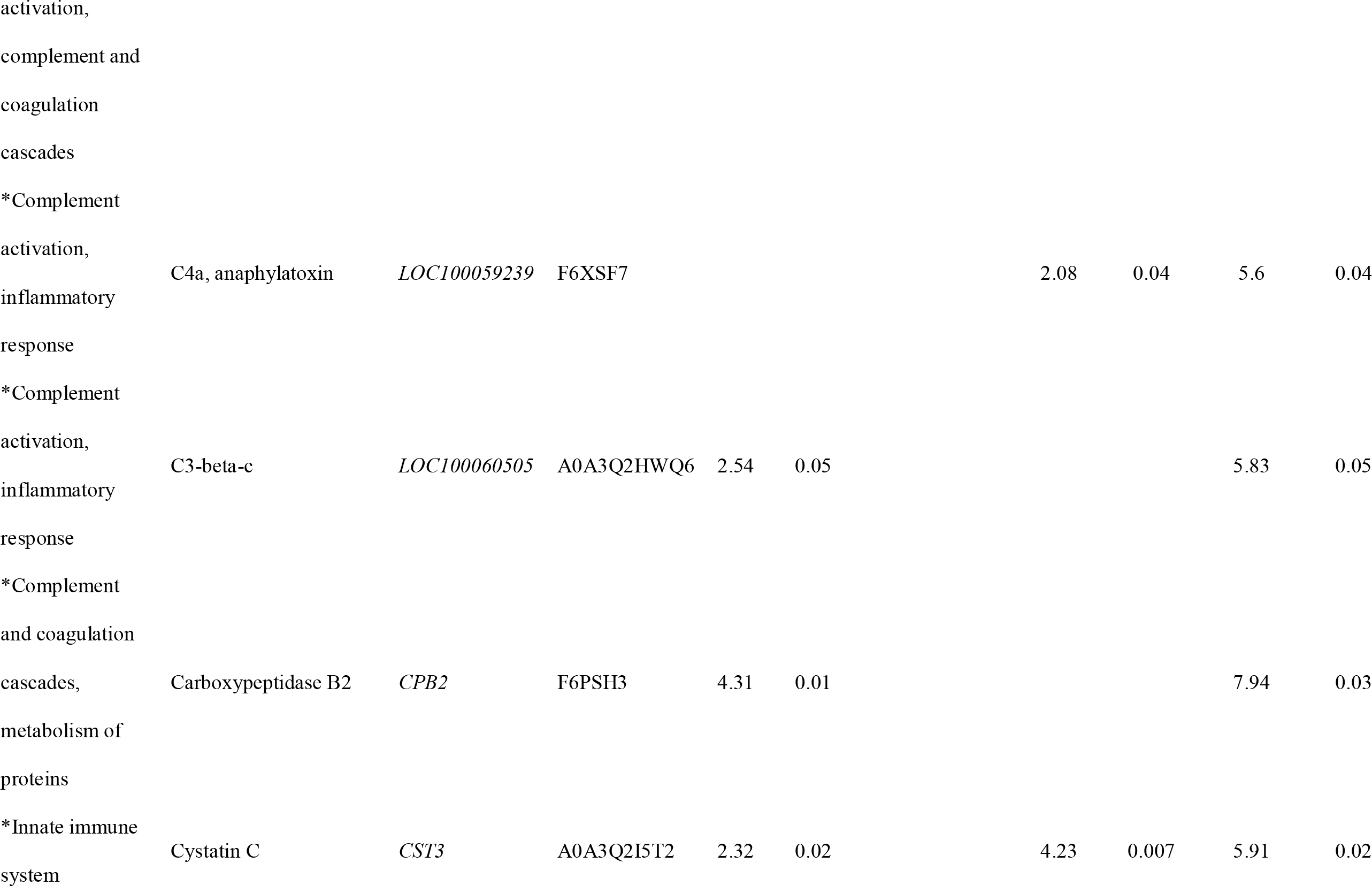
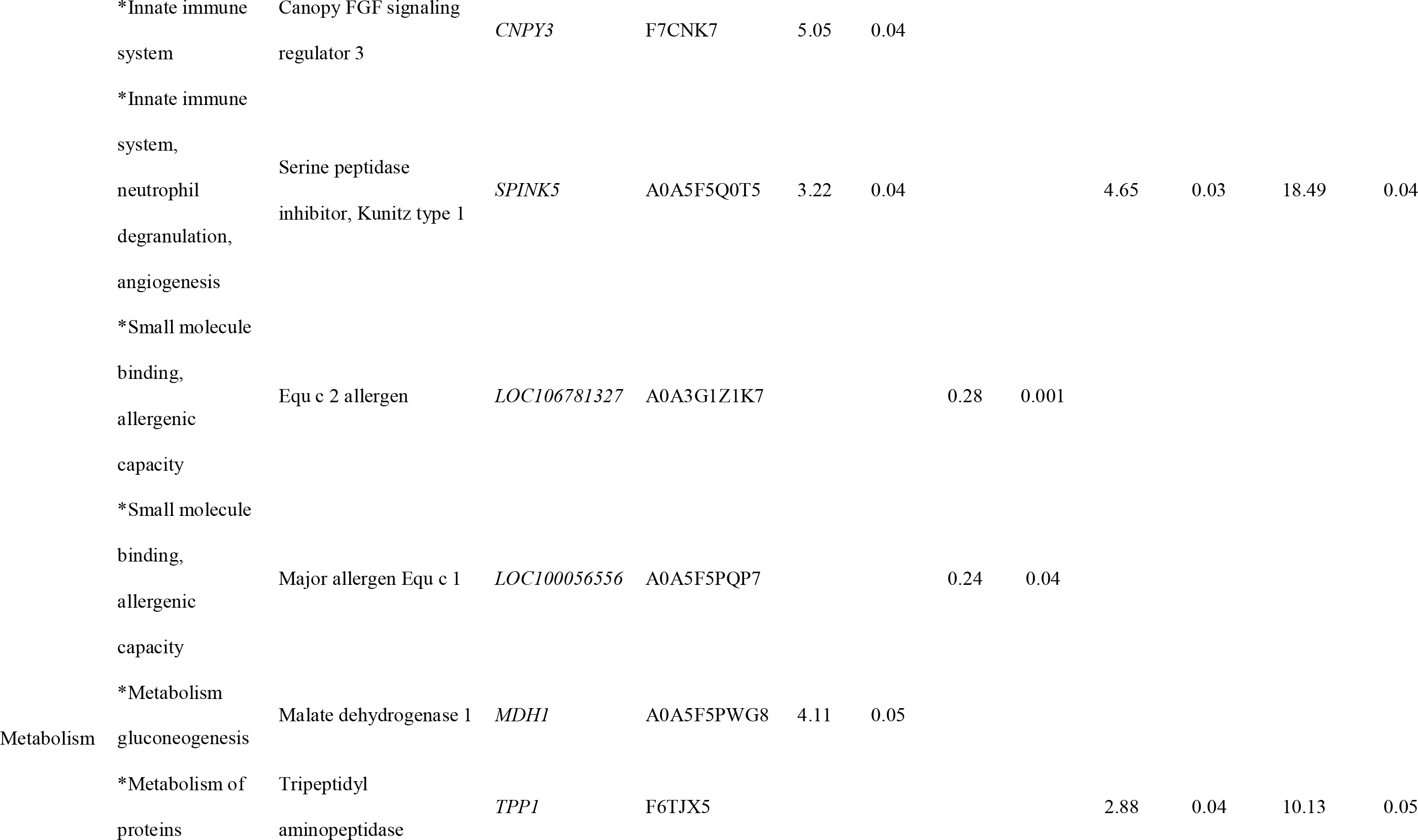
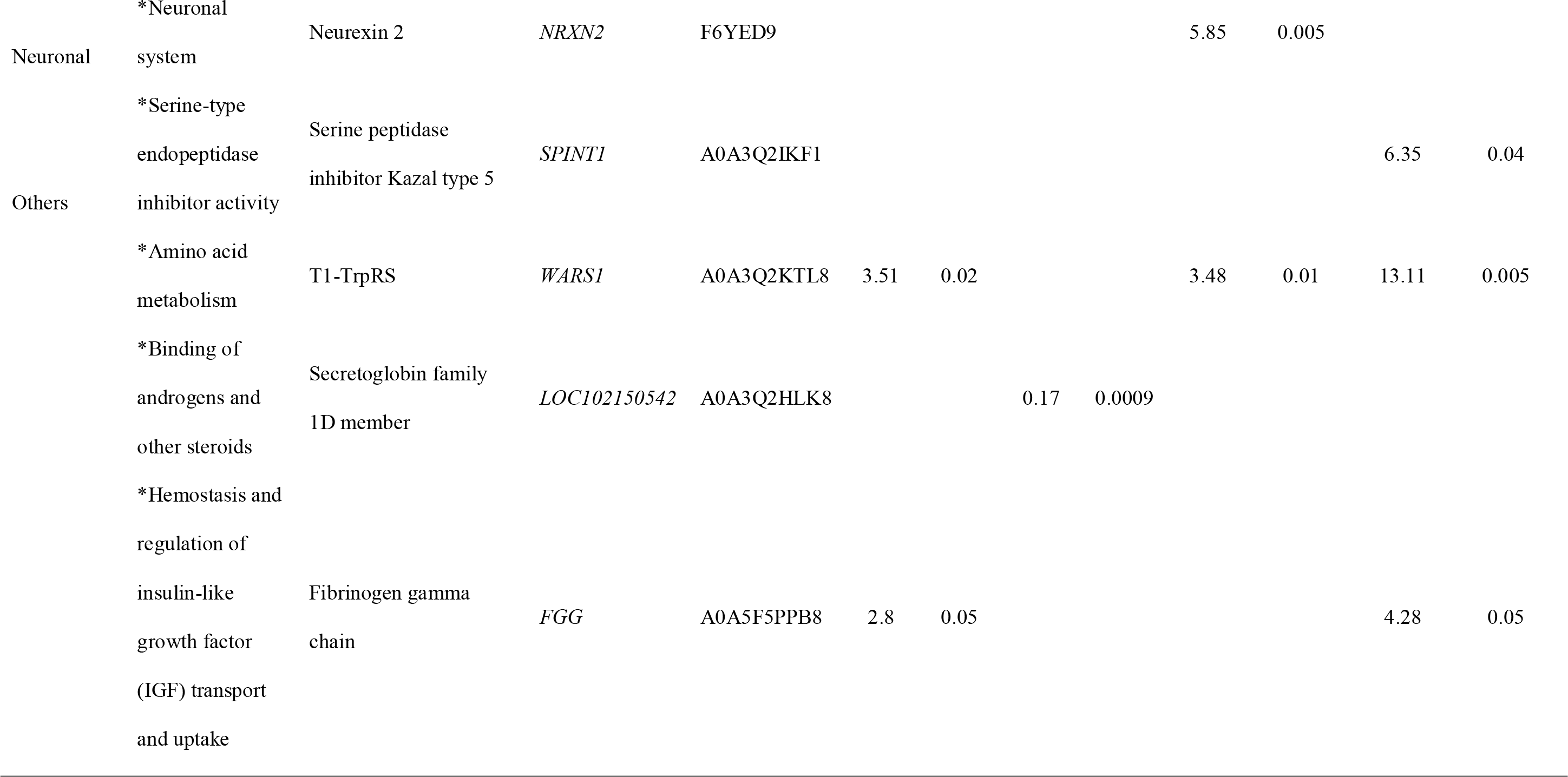
Fold-change and P-value for the top most upregulated proteins in each comparison in the quantitative proteomic analysis, including all teins significant for more than one comparison

### Place for Table 7, see end of document

When comparing protein levels during exercise vs. at rest (comparisons 1 and 2), there were 21 proteins in the SPH group and three proteins in the EPH group with significantly different levels (Supplementary Table S8). A gene ontology (GO) analysis for the 21 proteins showing changing expression profiles at rest vs. during exercise in the SPH group revealed pathways related to blood coagulation, hemostasis, coagulation and regulation of body fluid levels (Figure 5A) (32). When comparing EPH and SPH at rest (comparison 3), there were proteins upregulated in the SPH group (Supplementary Table S8), implicated in pathways related to immune response and complement activation (Figure 5B). There were no statistically significant differences in protein levels when comparing EPH and SPH during exercise (comparison 4). The paired ratio comparison (comparison 5) revealed 19 proteins showing significant difference in the individual difference at rest and exercise in SPH and EPH groups (Supplementary Table S8). These proteins are involved in pathways related to immune response, complement activation and catalytic and hydrolase activity (Figure 5C).

**Figure 5.**
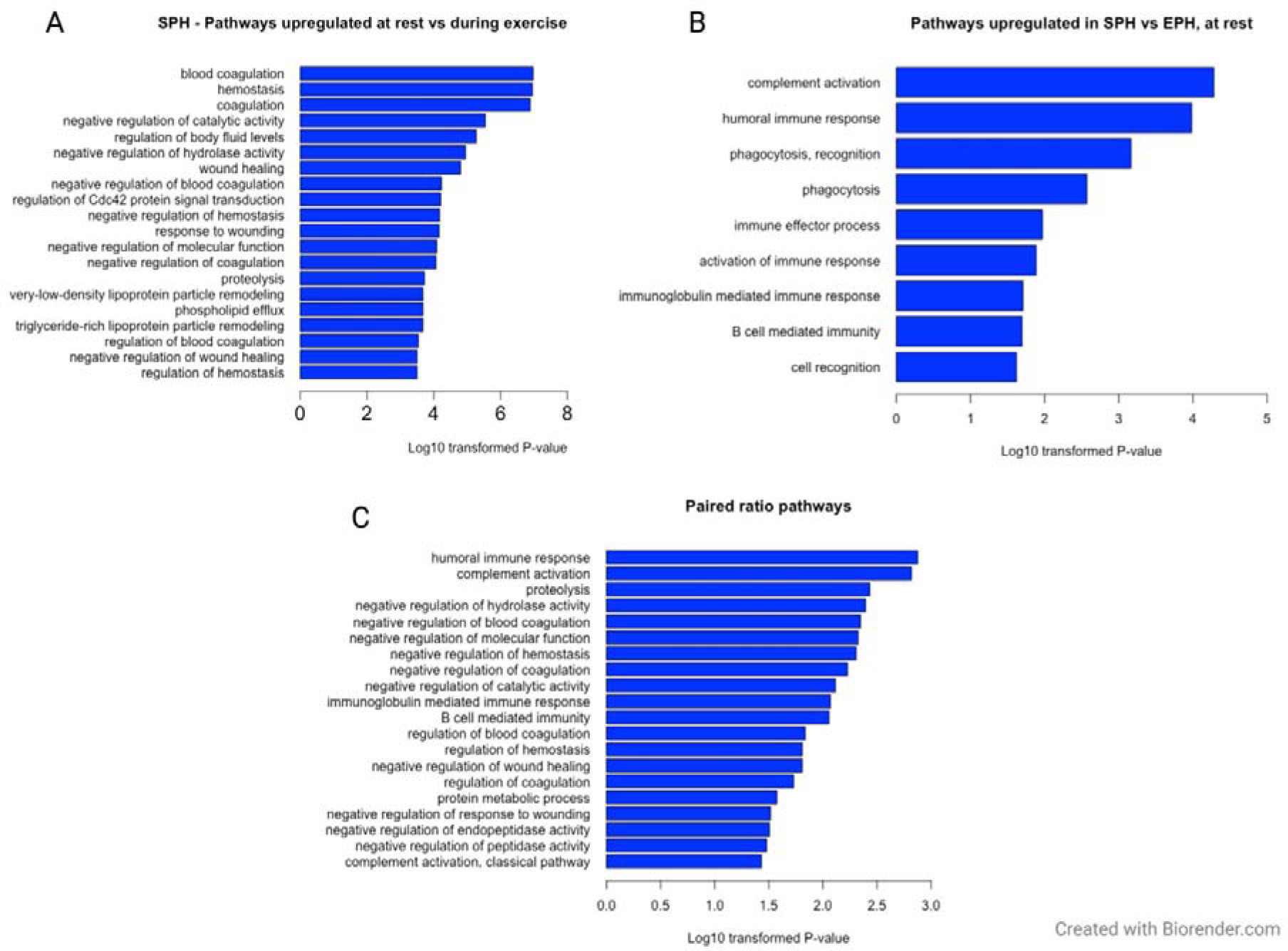
A. The 20 most significant pathways from the GO analysis based on the significant proteins in the comparison of exercise vs. rest in the SPH (sub-elite performing haplotype) group. B. All significant pathways from the GO analysis based on the significant proteins in the comparison of SPH (sub-elite performing haplotype) and EPH (elite performing haplotype) at rest. C. The 20 most significant pathways from the GO analysis based on the significant proteins in the paired ratio analysis of SPH vs. EPH.

## Discussion

The EDN3 protein has an established functional role in blood pressure regulation in humans and other species. However, to the best of our knowledge, there are no published reports establishing associations between the *EDN3* gene and equine athletic performance, except for our previous selective sweep study (13). Given the role of EDN3 in blood pressure regulation, we hypothesized that 1) an underlying phenotype for elite athletic performance is blood pressure regulation, and 2) the identified non-coding region harbors a regulatory element that acts on the *EDN3* gene.

All the SNPs defining the haplotypes were found to segregate in the genome of horses that lived prior to the rise and spread of the modern domestic lineage around ∼4,200 years ago (DOM2) (31). In fact, temporal allelic trajectories portray an increase in the frequency of the alternative allele, from 7,500-5,500 years ago, prior to the earliest archaeological evidence of horse husbandry (30, 33). The concerted rise in frequency of all alternate alleles following the spread of the DOM2 lineage is compatible with a haplotype undergoing positive selection, possibly following artificial selection for animals showing improved athletic performance. Interestingly, the allele trajectories at *EDN3* are in striking contrast with those previously described at the *MSTN* locus, driving performance in short-distance racing (6), which indicated selection starting within approximately the last 1,000 years only. This may indicate performance traits representing recurrent selection targets in the history of horse domestication.

We next demonstrated significant differences in blood pressure values, as well as protein levels of EDN1 and EDN3, when contrasting homozygous EPH to SPH CBT carriers. The 5.5 kb region contained multiple marks pointing at a potential *cis*-regulatory module. First, the region spanned an equine enhancer (H3K4me1) active in tissues relevant to racing performance. Second, comparative evidence from human gTEX v8 implicated this region in the regulation of *EDN3* expression across multiple tissues. Third, of the 14 SNPs forming the haplotypes, alleles from 12 showed differential transcription factor binding affinities, suggesting differential regulatory activity between the EPH and SPH carriers. Intriguingly, the SPH haplotype was predicted to have binding affinities for both immune system actors, NFATC2 and IRF2, where IRF2 has been shown to be a negative regulator of NFAT target genes (34) (Supplementary Table S4). On the other hand, the EPH haplotype harbored motifs for factors involved during embryogenesis and cell differentiation, e.g., SRF, HOXA5 and CREB1 (35–37) (Supplementary Table S4). These differing regulatory roles may point to the multifaceted role of EDN3, from early neural crest development and proper function of enteric neurons and melanoblasts (38). The EDN3 ligand has a broad expression range across tissues and has been implicated in diseases such as Hirschsprung’s disease, Multiple Sclerosis and Waardenburg syndrome in humans, as well as melanocyte development in mice (39–43). However, a key role for this ligand is in blood flow homeostasis.

Endothelins are involved in one of the most potent vasoregulatory systems, where EDN1 and EDN3 have opposite effects and act synergistically to regulate blood pressure (18). EDN1 increases blood pressure by vasoconstriction, while EDN3 is involved in nitric oxide release, which results in vasodilation and decreased blood pressure (18). Additionally, EDN3 stimulates the secretion of vasopressin, which increases the blood volume by retaining water in the kidneys, while EDN1 has the opposite effect in the kidneys (44–47). In the present study, horses homozygous for EPH showed significantly higher plasma concentrations of EDN3 and lower blood pressure during exercise. In contrast, horses homozygous for SPH had a significantly higher plasma concentration of EDN1 and higher blood pressure during exercise. Blood pressure depends on cardiac output and resistance in peripheral blood vessels, while total peripheral resistance is affected by the diameter of the arteries and the viscosity of the blood. An increase of EDN3 likely leads to a decrease in blood pressure and an increase in vasodilation of the blood vessels, primarily in the working muscles and the lungs. This means that blood can flow more easily, allowing improved arterial and venous oxygenation. This would, in turn, be beneficial for athletic performance. In humans, infusion of EDN3 caused a decrease in mean arterial blood pressure and vasoconstriction to the visceral organs (48). On the other hand, infusion of EDN1 leads to an increased mean arterial pressure and decreased cardiac output (49, 50). This is comparable to a study on horses during anesthesia, which revealed that horses receiving nitric oxide, a vasodilator, had better arterial and venous oxygenation and lower EDN1 plasma concentration than the control group without nitric oxide administration (51). In our study, we did not see any effect of exercise on the endothelin levels, as there were no statistically significant differences in EDN1 or EDN3 levels when contrasting resting and exercising. This is in concordance with previous studies in which no differences in EDN1 levels before and during exercise were observed in healthy horses (22, 52). In terms of exercise performance, the present study showed that SPH homozygous horses had a slower recovery than the horses carrying the EPH. This could be because the SPH group had higher blood pressure during the exercise, so it took longer for these individuals to return to resting blood pressure values.

Although the current study demonstrated statistically significant differences in EDN1 and EDN3 levels between the different haplotype groups, there were no differences in blood pressure at rest. In adult horses, normal blood pressure at rest is approximately 130/95 mmHg (systolic/diastolic), in line with the values observed in the current study (53). In humans, it has been demonstrated that elevated blood pressure at rest is negatively correlated with athletic performance (21). Individuals with elevated blood pressure had significantly lower maximal oxygen consumption, ventilatory anaerobic thresholds and heart rate reserves (difference between an individual’s resting heart rate and maximum heart rate) (21). However, unlike humans, high blood pressure in horses is uncommon and most blood pressure measurements on horses are performed to evaluate and monitor hypotension. An increase in blood pressure at rest is most commonly seen as a result of diseases such as laminitis, chronic renal failure or equine metabolic syndrome (54–56).

The spleen represents one important organ for the distribution of blood, acting as a reservoir for red blood cells. The equine spleen reservoir is larger than in any other domestic animal (57). When there is an increased demand, red blood cells stored in the spleen can be released into the system (57). During exercise, the spleen is contracted, which causes an increase in the volume of circulating blood cells, while the volume of plasma is unchanged or even reduced. This leads to an increase in the packed cell volume/hematocrit (i.e., the proportion of blood made up of red blood cells), hemoglobin concentration, and red blood cell count (57). Previous studies have demonstrated significantly larger spleen size in horses used for racing compared to breeds that are not traditionally used for racing (58). In addition, one study demonstrated a significant correlation between blood pressure and spleen volume (59), with the splenic volume decreasing when hypertension was induced. However, when hypotension was induced, there were no significant changes in the splenic volume (59). The results by (59) demonstrated the importance of the spleen for the cardiovascular system in horses. Further studies examining the relationship between blood pressure, spleen volume and expression of EDN3 represent promising avenues for enhancing understanding of the cardiovascular system in horses in relation to exercising.

Our global proteomic analysis resulted in the identification of a total of 582 proteins. Only one such protein, Cathepsin X, was encoded as part of the 20 protein-coding genes located within 2 Mb surrounding the genomic region investigated (ECA22: 45,718,095 – 47,718,095). In contrast to the ELISA analysis, none of the endothelins were detected. A major obstacle in MS-based plasma proteomics is the large dynamic range in protein concentrations and that the abundant high molecular weight proteins mask the identification and quantification of lower molecular weight proteins, such as the endothelins. The highly abundant proteins contribute to approximately 90 % of the total protein content in plasma. In this study, plasma depletion of the most abundant proteins was performed to facilitate the quantification of less abundant proteins and to improve the sequence coverage (60). However, a limitation of the present study is the quality of the available proteome database for *Equus caballus*, which only includes a limited number of proteins. Additionally, many proteins were excluded due to a low number of biological replicates, and in total, 383 proteins could be included in the comparisons between groups.

When comparing protein levels at rest and during exercise for the different haplotype groups, we observed a clear difference in the number of regulated proteins (Supplementary Table S8). In the SPH group, 21 proteins were detected at higher levels at rest, than during exercise. The GO analysis of these proteins identified several pathways related to injuries, including blood coagulation, hemostasis, and wound healing (Figure 5A). Several of these proteins also have functions related to the immune system. When comparing the protein levels between the haplotype groups at rest, 23 proteins were found regulated. The GO analysis revealed significant pathways mainly related to the immune system. This is consistent with previous studies in which an increase of plasma proteins involved in pathways related to inflammation, coagulation and immune modulation was observed (61, 62). A previous genome-wide association analysis of racing performance in CBTs also found correlations for genes related to immune response (63). Racing horses perform intense exercise, which induces a physical stress response to obtain energy for the working muscles. The released stress hormones (e.g., cortisol) stimulate gluconeogenesis and lipolysis (64). Additionally, the stress hormones affect inflammatory and coagulation processes (61,62,65,66). In a non-inflammatory state, intact endothelial cells maintain blood flow and control coagulation. However, chronic inflammation, oxidative stress and mechanical stress from turbulent blood flow negatively affect the function of the endothelial cells (67, 68). Furthermore, in a human study, multi-omics changes in response to exercise were observed, involving pathways related to energy metabolism, oxidative stress and immune response (69). Potentially, SPH horses have more difficulties recovering from exercise due to elevated blood pressure, resulting in higher levels of immunity-related proteins and proteins related to injury healing in the plasma at rest.

It is interesting to note that the protein showing the highest expression difference between SPH and EPH groups at rest was the proteasome subunit beta (PSMB10). Interestingly, the *psmb10* gene has previously been linked to blood pressure regulation in mice. Knockout of the *psmb10* gene was protective for high blood pressure, cardiac fibrosis and inflammation in DOCA salt-treated mice (70). Although the *PSMB10* gene may be involved in blood pressure regulation in the horse, it is notable that there were no differences in blood pressure between the haplotype groups at rest. In the EPH group comparison of protein ratios at rest versus exercise, there were only three proteins with significant differences. Notably, one of the proteins detected was the secretoglobin family 1D member (LOC102150542). Secretoglobins are known to have anti-inflammatory effects in airway disorders such as asthma in both horses and humans (71, 72). The increased levels of this protein may play an anti-inflammatory role during exercise by preserving the function of the vascular endothelial cells, contributing to a higher tolerance for intense exercise.

Several studies have demonstrated the effect of the *EDN1* gene and the endothelin receptor gene, *EDNRB*, in horses and humans (73–81). Studies have suggested EDN1 as a marker for different diseases in horses, for example, hyperinsulinemia, neonatal diseases, and cardiac disease (76–78). Moreover, the EDNRB receptor is known to be associated with lethal white foal syndrome (73–75). However, to the best of our knowledge, there is only one study performed on horses involving the *EDN3* gene (13). The current study demonstrates significant differences in blood pressure and plasma levels of both EDN3 and EDN1 in CBTs with different haplotypes. Although the identified gene variant may also act on other genes in the region or other parts of the genome, we believe that we have identified a regulatory variant influencing the transcription of *EDN3*. Based on the ELISA results and racing performance data, the identified candidate SNP appears to have an additive effect, with homozygous carriers and heterozygous horses performing significantly better than homozygous wild-type horses. The obtained results in horses emphasize the importance of the endothelin system as critical for performance and support the functional role previously shown for EDN3 in human blood pressure regulation.

## Conclusion

Here we demonstrate, for the first time, a significant association between *EDN3*, blood pressure regulation and athletic performance in horses. We identified a 5.5 kb candidate genomic interval and showed that horses homozygous for the elite performing haplotype had a higher plasma concentration of EDN3, lower plasma concentration of EDN1, and lower blood pressure during exercise compared to horses with the sub-elite performing haplotype. Horses carrying the elite performing haplotype also recovered from exercise faster than those with the sub-elite haplotype. The results reported here are important for understanding the biological mechanisms behind blood pressure regulation, as well as inflammation and coagulation systems in relation to racing performance in both horses and other mammals, including humans.

## Materials and Methods

### Horse material and DNA extraction

A summary of the horses included in the study, and specifying which experiments they were used for, is presented in Supplementary Table S2. Genomic DNA from all horses was extracted from hair roots or blood samples. For hair preparation, 186 μL of 5 % Chelex® 100 Resin (Bio-Rad Laboratories, Hercules, CA, USA) and 14 μL of proteinase K (20 mg/mL; Merck KgaA, Darmstadt, Germany) were added to each sample. The mixture was incubated at 56°C for two hours at 600 rpm and inactivated with proteinase K for 10 mins at 95°C. Blood samples were extracted on the Qiasymphony instrument with the Qiasymphony DSP DNA mini or midi kit (Qiagen, Hilden, Germany).

### Selective sweep racing performance association analysis with additional horse material

We used the previously identified 19.6 kb selective sweep region, associated with CBT racing performance, as the starting point for this analysis (ECA22:46,702,297-46,721,892; EquCab3.0) (13). In that study, g.46717860-T was the variant most significantly associated with elite performance, and LD to this variant was used to define a five SNP haplotype (*r^2^* > 0.9). We extracted the 19.6 kb region from an existing 670K Axiom Equine Array genotyping experiment of 661 CBTs (including the 400 horses used for the haplotype analysis in the 2018 study) (13, 63). Using g.46717860 as the lead variant, we calculated pairwise r^2^ across the region, and performed racing performance association analyses between this variant, and any additional variant with r^2^ > 0.6. Given the high LD, a significance threshold was set at P ≤ 0.05.

The available phenotypic records included a) pedigree information b) *DMRT3* genotypes and c) performance data as individual race records for each horse (Swedish Trotting Association (Svensk Travsport) and the Norwegian Trotting Association (Norsk Rikstoto). The *DMRT3* variant was genotyped using StepOnePlus Real-Time PCR System (Thermo Fisher Scientific, Waltham, MA, USA) with the custom TaqMan SNP genotyping assay (82), and included as a factor in the model, since previous studies have demonstrated major effects of the variant on harness racing performance (10, 12). Only competitive races were included (i.e., premie and qualification races were excluded). The performance data included: number of starts, number of wins, number of placings (the number of times a horse finished a race in first, second or third place), fastest kilometer (km) time in a race where the horse did not gallop (in seconds), earnings and earnings per start. The earnings for most horses were recorded in Swedish currency (SEK). Earned prize money in Norwegian currency was converted to SEK based on the average exchange rate for the year the race took place.

All statistical analyses were performed in the R statistical environment. The performance data were tested for normality by computing the skewness coefficient using the package moments v0.14. Non-normally distributed values were transformed, i.e., log transformed values (log10 +1) were used for wins and placings, number of starts were square root transformed and earnings and earnings per start were transformed according to the previously reported formula (ln(earnings + 1000)) (83). Genetic variants were tested in a linear model using ANOVA as a *post hoc* test. Number of starts, age at first start, sex, birth year, country of birth, and the *DMRT3* genotype were included as fixed effects, when significant. Significant values (P ≤ 0.05) were further tested using the function lsmeans (Least-Squares Means) with the package emmeans followed by the multiple comparison test Tukey’s HSD-test.

### Additional variant discovery

To find and assess variants missing from the Axiom Equine Array, but with the potential to influence racing performance, we performed both Illumina short-read WGS and MinION Oxford Nanopore targeted sequencing of PCR products.

First, two CBTs and two SBs were selected for WGS based on their g.46717860 genotype and their performance values. From each breed, horses homozygous for opposite alleles of SNP g.46717860 (CC respectively TT) with either high (TT) or low (CC) earnings per start were selected for sequencing (Supplementary Table S4). Paired-end 150 bp Illumina HiSeqX data was generated to a depth of 15X. Read data were processed according to the GATK 3.8- 0 best practices (https://software.broadinstitute.org). Briefly, raw data were trimmed with TrimGalore 0.4.4 and quality checked with QualiMap 2.2. The reads were aligned to the EquCab3.0 reference genome (84) (GCF_002863925.1) using BWA-MEM 0.7.17. SAM files were converted to BAM files using SAMtools, and Picard 2.10.3 was used to sort the BAM files and remove potential PCR duplications. The BAM files were recalibrated in two steps using BaseRecalibrator and PrintReads. Variant calling was performed using HaplotypeCaller with a list of known variants in horses downloaded from the NCBI web page annotation release 103 (https://www.ncbi.nlm.nih.gov). Structural variants were called using default parameters in FindSV comprising the software programs Manta, TIDDIT and CNVNator (https://github.com/J35P312/FindSV).

Next, as input for Oxford Nanopore sequencing (ONT, Oxford, UK), a 25 kb PCR product was designed to span the 19.6 kb sweep region and adjacent repeats. Primers were designed in Primer3Plus and included unique 10 bp barcodes (Supplementary Table S9). DNA input was taken from four CBTs with g.46717860-TT genotype and high performance and four CBTs with the opposite features, including the two CBTs from the WGS analysis (horse selection as per WGS above and Supplementary Table S2). The PCR region was amplified on a ProFlex PCR cycler (Thermo Fisher Scientific, Waltham, MA, USA) and the long-range PCR using AccuPrime™ Taq DNA Polymerase, High Fidelity kit following manufacturers specifications. The sizes of PCR products were confirmed on 0.8 % TBE Agarose gels, and product concentration quantified with Agilent 4150 TapeStation using the Agilent Genomic DNA ScreenTape Assay (Agilent Technologies, Santa Clara, CA, USA). Samples of the respective PCR products were pooled in equimolar concentration and libraries were prepared using the Genomic DNA by Ligation (SQK-LSK109) protocol. The samples were loaded and sequenced on the MinION. Demultiplexed sequencing data were mapped to EquCab3.0 ECA 22 (GCF_002863925) using minimap2 v. 2.4 sequence alignment program (https://github.com/lh3/minimap2). The resulting SAM files were converted to BAM format using samtools v. 1.8. Single nucleotide variants were called using the variant calling tool longshot v. 0.4.3 with default parameters (https://github.com/pjedge/longshot). Structural genomic variations were identified with NanoSV v. 1.2.4 software (https://github.com/mroosmalen/nanosv), with default parameters. Only variants with a quality above 20 were retained.

### Genotyping

The largest INDEL identified from the WGS data was analyzed for association with racing performance traits in 369 CBTs, using linear models. For MassArray SNP genotyping the number of variant positions available was limited to 25. For this reason, 24 SNPs and one INDEL, evenly distributed across the 19.6 kb region, were selected for genotyping. In total, DNA from 412 horses of 13 different breeds and four donkeys were genotyped (Supplementary Table S10). The horses were carefully selected to include both traditional performance horse breeds, as well as non-performing breeds. Genotyping was performed at the Mutation Analysis Facility at Karolinska University Hospital (Huddinge, Sweden), using iPLEX® Gold chemistry and the MassARRAY® mass spectrometry system (85) (Agena Bioscience, San Diego, CA, USA). Multiplexed assays were designed using MassARRAY® Assay Design Suite v2.2 software (Agena Bioscience, San Diego, CA, USA), genotyping the 25 markers in one reaction per sample. The protocol for allele-specific base extension was performed according to Agena Bioscience’s recommendation. Analytes were spotted onto a 384-element SpectroCHIP II array (Agena Bioscience, San Diego, CA, USA), using Nanodispenser RS1000 and subsequently analyzed by MALDI-TOF on a MassARRAY® Analyzer 4 mass spectrometer (Agena Bioscience, San Diego, CA, USA). Genotype calls were manually checked by two persons individually using MassARRAY® TYPER v4.0 Software (Agena Bioscience, San Diego, CA, USA).

The variant g.46717860 was not included as part of the MassArray, but was genotyped for 170 horses (9 Ardennes, 96 CBTs, 4 donkeys, 4 Finnhorses, 2 Fjord horses, 2 Gotlandsruss, 10 North-Swedish draught horses, 3 Pzrewalski and 40 SBs) using the StepOnePlus Real-Time PCR System (Thermo Fisher Scientific, Waltham, MA, USA) and the custom TaqMan SNP genotyping assay (82). The reaction volume of 15 µl comprised 1.5 µl DNA, 0.38 µl Genotyping Assay 40X, 7.5 µl Genotyping Master Mix 2X, and 5.62 µl deionized water.

### Haplotype analysis

Pairwise r^2^ calculations with all 24 SNPs and g.46717860 were performed using the LD- function in R. Based on the high LD between all genotyped SNPs, it was possible to impute g.46717860 alleles for ungenotyped samples. All SNPs with a pairwise r^2^ value >0.6, plus two flanking SNPs were selected for haplotype analysis. Haplotype racing performance association tests used a GLM regression analysis, which was performed in 180 CBTs and 38 SBs using the haplo.glm function from the haplo.stats package in R (86). The model included the effects of sex, age, country of registration, *DMRT3* genotype (only for the CBTs) and number of starts, when significant. All CBTs had been genotyped for the DMRT3 mutation, using the StepOnePlus Real-Time PCR System (Thermo Fisher Scientific, Waltham, MA, USA) with the custom TaqMan SNP genotyping assay (82). Haplotypes with a frequency < 2 % were considered rare, and were not analyzed for association with performance. A minimum shared haplotype of 14 SNPs between CBTs and SBs was used to define the elite- performing haplotype (EPH) and the sub-elite performing haplotype (SPH).

### Association analysis with athletic performance

Allele calls at SNP g.46718095 were a proxy for elite-performing haplotype (EPH) and the sub-elite performing haplotype (SPH). Samples not genotyped on the MassArray were genotyped using the StepOnePlus Real-Time PCR System (Thermo Fisher Scientific, Waltham, MA, USA) with the custom TaqMan SNP genotyping assay (82). Association with racing performance was performed as per “Selective sweep racing performance association analysis with additional horse material” using 539 CBTs born between 1994 and 2017 and 271 SBs born between 2005 and 2014. Horses of 13 different breeds and three other types of equids (wild ass, onager and donkey) were also genotyped to investigate the distribution of allele frequencies.

### Functional bioinformatic analyses

ChIP-seq data were provided by the Equine Functional Annotation of Animal Genomes (FAANG) team (www.faang.org) (24, 25) including four histone marks (H3K4me1, H3K4me3, H3K27ac and H3K27me) across nine tissues (adipose, brain, heart, lamina, liver, lung, muscle, ovary and skin) from two Thoroughbreds. FAANG bam and bed files from the different available tissues, the four histone marks and the input DNA control (project: PRJEB35307) were downloaded to a local server. First, the integrity of the data was checked using checksums. Bam files were sorted with samtools and bedgraphs were subsequently created using bedtools (87, 88). The Integrative Genomics Viewer (IGV) browser was used to visualize the output data along with the EquCab3.0 fasta file and its gff3 annotation file (84, 89). The presence of H3K27ac and H3K4me1 was considered to mark active enhancers, whereas H3K4me1 and H3K27me3 marked poised enhancers. H3K27ac and H3K4me3 marked active promoters and H3K27me3 was considered to mark repressed regions. The region defined by the minimum shared haplotype (see above) was investigated using IGV for the different histone marks in all available tissues.

A comparative analysis to hg38 was conducted for the minimum shared haplotype region. Co-ordinates were lifted in UCSC (liftOver) and potential overlaps between ENCODE Candidate Cis-Regulatory Elements (cCREs), ENCODE Integrated Regulation tracks and ORegAnno Regulatory elements were annotated (90, 91). For each of the SNPs in the minimum shared haplotype region, the potential of alleles to alter transcription factor binding potential was tested with sTRAP (92). Here, 31 bp sequences centered on the allele of interest were tested using JASPAR vertebrates, on a background model of human promoters. In the cases where two SNPs were within 10 bp of each other, they were tested as a single phased fragment.

### Ancient DNA screening

We leveraged the availability of an extensive ancient genome time-series in the horse to assess the temporal trajectory and spatial distribution of the SNPs in the minimum shared haplotype within the CBTs, in the past. More specifically, we re-aligned against the EquCab3.0 reference genome (84) supplemented with the Y-chromosomal contigs from (93) and a sub-selection of the sequencing data from (2,27,29,30,94,95), representing a total of 431 ancient genomes. Alignment files were generated using the Paleomix pipeline (96), and further rescaled and trimmed, following the procedures described in (31). This procedure ensured minimal sequencing error rates, especially at sites potentially affected by post-mortem DNA damage. The number of occurrences of each individual allele was counted for each individual using ANGSD (v0.933), with the -baq 0, -remove_bads -minMapQ 25 -minQ 30 -rmTriallelic 1e-4 -SNP_pval 1 -C 50 parameters (97). The resulting occurrences were tabulated together with metadata available for each individual, consisting of their corresponding archaeological site (name and GPS coordinates) and their average calibrated radiocarbon date (or age as assessed from the archaeological context otherwise). The tabulated file was visualized using the mapDATAge package (26), which provides spatial distributions within user-defined time periods and/or spatial ranges and estimates allelic trajectories (i.e., allelic frequencies through time) based on the resampling of individual allelic counts. The final allelic trajectories were assessed considering time bins of 2,000 years (step-size = 500). To explore the genetic variation for the SNPs within the haplotype, we considered the 14 polymorphic sites i.e., ECA22:46703380, 46708110, 46708983, 46713478, 46715974, 46717451, 46717528, 46717742, 46717854, 46717999, 46718095, 46718113, 46718361, 46718436, 46718895, 46718964, 46719042, 46719906, 46720030, 46720999, and 46721822.

### Exercise test, sample collection and blood pressure measurements

#### Horse material

Thirty horses were enrolled in the exercise test (Supplementary Table S2). All horses were genotyped for SNP g.46717860 and g.46718095 using the StepOnePlus Real-Time PCR System (Thermo Fisher Scientific, Waltham, MA, USA) with the custom TaqMan SNP genotyping assay as above (82). The two SNPs were used as a proxy for the haplotype variants homozygote EPH (TT respectively CC) or homozygote SPH (CC respectively TT). Horses were then divided into three groups based on haplotype variant, i.e., homozygote EPH or SPH and HET (heterozygote). The horses were carefully selected to include an even sex and age distribution across the different groups. The age of the horses ranged from two to 13 years (mean age = 5.5 years). All horses, except for the two-year old, were in competing condition. However, due to various reasons including practical problems, blood pressure measurements were not taken from all horses at all time points and some horses only had measurements at rest (Supplementary Table S2). For the ELISA and proteomics analysis, plasma samples were collected from 40 horses, before and/or during exercise. As for blood pressure measurement, some horses only had samples collected at rest.

#### Exercise

All horses, except for the two two-year old horses, were challenged on the same standardized exercise session (n=28). The test was performed on several days during autumn and winter of the years 2020 and 2021. Each horse, with a trainer, completed a 1.5-hour exercise route that included both flat and uphill interval training. This route was commonly used in their daily training program. A Polar M460 sensor with GPS (Polar Electro, Kempele, Finland) was used to record heart rate (HR), speed, distance and amplitude during exercise. The horses performed the exercise on a track with hilly terrain in the forest. The uphill interval section consisted of the horses trotting four times up a hill with four degrees inclination. Each uphill interval lasted for five mins with a distance of 1.7 km. The HR for each horse reached above 200 beats/min. After the uphill intervals, the horses walked down to the flat part of the track, where they completed 10 mins of intervals, at a speed of 35 km/h and HR of about 180-200 beats/min. Finally, the horses jogged one km back to the stable at a speed of 15-20 km/h. During the exercise, the average HR reached above 190 beats/min for approximately eight mins. Due to their young age the two-year old horses (n=2) performed a standardized exercise session at the racetrack. Following warm-up, they performed a number of intervals with HR reaching over 200 beats/min. The two-year old horses performed the exercise test at the same time.

#### Blood pressure measurements

The following measurements were collected with a Cardell Veterinary Monitor 9402 cuff (Cardell 10), at the middle coccygeal artery: systolic, diastolic, mean arterial pressure, pulse pressure, and heart rate. The same cuff size was used for all the horses. The monitor has previously been successfully evaluated for use in horses (98). Blood pressure was measured at rest in the stable before the exercise and during exercise (directly and five mins after the last uphill interval) (Figure 6). Additionally, blood pressure was measured every ten mins after exercise, up to 40 mins after the exercise was completed. In order for the horse to acclimate to the equipment, the Polar M460 sensor and the cuff were put on at rest, before the exercise route, and the horse was left in the box for 10 mins before measurements were taken. Similarly, blood pressure was measured five times at rest for the horse to get used to the measurement procedure. The measurements with the highest and lowest systolic blood pressure values were removed. During and after exercise the blood pressure was measured three times and an average calculated. For the association analyses with blood pressure, the mean values of the triplicates were analyzed using ANOVA and Tukey’s HSD test.

**Figure 6.**
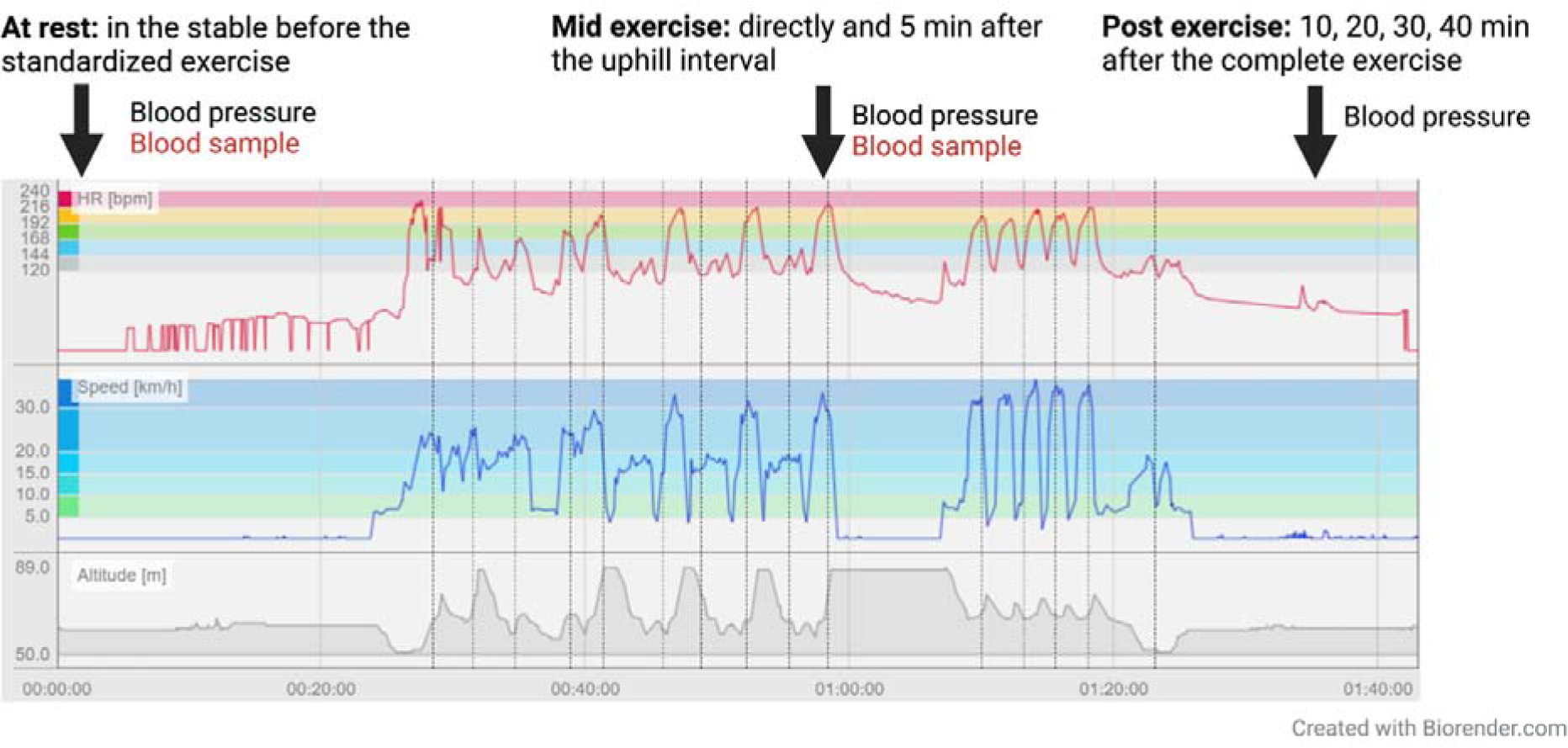
A schematic overview of the parameters recorded during the exercise for each horse, including heart rate, speed and altitude. The arrows indicate when blood samples and blood pressure measurements were taken.

#### Biological sample collection and analysis

Blood samples were collected at rest before the exercise and during exercise directly after the last uphill interval. Blood was also collected for plasma extraction. Here, a sample was collected in an EDTA tube and mixed gently four times after sampling, to allow the EDTA to mix with the blood, then centrifuged at 1000 x g (3000 rpm) for 20 mins as soon as possible and within two hours after sampling. The tubes were kept cold until centrifugation. The EDTA plasma was aliquoted in Eppendorf tubes and immediately frozen at -20°C and within five days put at -80°C until analysis.

### ELISA analysis of EDN1 and EDN3

Plasma EDN1 concentrations were measured using the Horse Endothelin ELISA Kit PicoKine™ EK0945-EQ (Boster Bio, Pleasanton, CA, USA). Plasma EDN3 was quantified with the horse EDN3 ELISA Kit MBS9901200 (MyBioSource, San Diego, CA, USA). The analyses were performed according to the manufacturer’s manual. A standard curve was created, and the samples were run in duplicates. Mean values of the concentration (pg/ml) were used in the statistical analysis, and the standard deviation (SD) and coefficient of variation (CV) were calculated. Differences in concentrations between the three groups (EPH, HET and SPH) before and during exercise were analyzed using ANOVA and Tukeýs HSD test. Significance was set at *P-*values ≤ 0.05.

### Relative quantitative proteomic analysis

A global relative quantitative proteomic analysis was performed to identify differentially expressed proteins in plasma before and during exercise. Six horses of each haplotype group (EPH or SPH homozygote) were included in the analyses (Supplementary Table S2). The horses were selected based on earnings per start, age and sex. The two-year-old horses were matched, one from each haplotype group, as they did not perform the same standardized exercise as the other horses.

### Sample preparation

The plasma samples and references (10 µl) were depleted from albumin and IgG, using the Pierce™ Albumin/IgG Removal Kit (Thermo Fisher Scientific, Waltham, MA, USA) according to the manufacturer’s instructions. The reference was a representative pool that included four groups: EPH at rest, EPH during exercise, SPH at rest and SPH during exercise. The samples were processed using the modified filter-aided sample preparation (FASP) method (99). Half the volume, corresponding to approximately 30 µg, was concentrated on Microcon-30kDa Centrifugal Filter Units (Merck & Co, Kenilworth, NJ, USA), reduced with 100 mM dithiothreitol at room temperature for two hours, washed several times with 8 M urea and once with digestion buffer (DB) (25 mM TEAB, 0.5 % sodium deoxycholate) before alkylation with 10 mM methyl methanethiosulfonate in digestion buffer for 30 mins. Digestions were performed by adding 1 µg Pierce MS grade Trypsin (Thermo Fisher Scientific, Waltham, MA, USA) in DB and incubating overnight at 37°C. An additional portion of 1 µg trypsin was added, and the samples were incubated for another three hours. Peptides were collected by centrifugation and labeled using Tandem Mass Tag (TMTpro 16plex) reagents (Thermo Fisher Scientific, Waltham, MA, USA), according to the manufacturer’s instructions. The labeled samples were combined into two sets, where both sets included all four groups and a reference. The paired samples were analyzed in the same set. Sodium deoxycholate was removed by acidification with 10 % TFA and the sets were desalted using Pierce peptide desalting spin columns (Thermo Fisher Scientific, Waltham, MA, USA) according to the manufacturer’s instructions. The two sets were pre-fractionated into 40 fractions with basic reversed-phase chromatography, using a Dionex Ultimate 3000 UPLC system (Thermo Fisher Scientific, Waltham, MA, USA), an XBridge BEH C18 column (3.5 μm, 3.0x150 mm, Waters Corp, Milford, MA, USA) and a gradient from 3 % to 90 % acetonitrile in 10 mM ammonium formate with pH 10.0 for 25 mins. The fractions were concatenated into 20 fractions, dried, and reconstituted in 3 % acetonitrile and 0.2 % formic acid.

### Liquid chromatography–Mass Spectrometry/ Mass Spectrometry analysis

The fractions were analyzed on an Orbitrap Fusion Lumos Tribrid mass spectrometer interfaced with an Easy-nLC 1200 liquid chromatography system (Thermo Fisher Scientific, Waltham, MA, USA). Peptides were trapped on an Acclaim Pepmap 100 C18 trap column (100 μm x 2 cm, particle size 5 μm, Thermo Fisher Scientific, Waltham, MA, USA) and separated on an analytical column (75 μm x 35 cm) packed in-house with Reprosil-Pur C18 material (particle size 3 μm, Dr. Maisch, Ammerbuch, Germany) at a flow of 300 nL/min using a gradient from 4 % to 80 % acetonitrile in 0.2 % Formic Acid over 90 mins. The Orbitrap was equipped with the FAIMS Pro ion mobility system alternating between the compensation voltages of -50 and -75. MS scans were performed at 120,000 resolutions, and the 12 most abundant doubly or multiply charged precursors were isolated using 0.7 m/z isolation window and dynamic exclusion within 10 ppm for 60 seconds. The precursors were fragmented by collision induced dissociation (35 %) and detected in the ion trap, followed by multinotch (simultaneous) isolation of the top 10 MS2 fragment ions within the m/z range 400-1400, MS3 fragmentation by higher-energy collision dissociation (55 %) and detection of the reporter ions in the Orbitrap at 50 000 resolution, m/z range 100-500.

### Proteomic data analysis

The data files in each set were merged for identification and relative quantification using Proteome Discoverer version 2.4 (Thermo Fisher Scientific, Waltham, MA, USA). The Uniprot *Equus caballus* proteome database (July 2021, combined SwissProt and TrEMBL 44,488 entries) and Mascot search engine v. 2.5.1 (Matrix Science, London, UK) were used in the database matching with peptide and fragment ion tolerance of 5 ppm and 0.6 Da. Tryptic peptides with zero missed cleavage, variable modifications of methionine oxidation and fixed cysteine alkylation were accepted. ProTMT modifications of N-terminal and lysine were selected. Percolator was used for the peptide-spectrum match validation with a strict false discovery rate (FDR) threshold of 1 %. The TMT reporter ions were identified with 3 mmu mass tolerance and no normalization was applied to the samples. For the protein quantification, only unique peptide sequences at a strict FDR threshold of 1 % with a minimum synchronous precursor selection match of 65 % and an average S/N above 10, were taken into account. The reference in each set was used to calculate protein ratios and quantified proteins were filtered at 5 % FDR.

### Processing of quantitative proteome data

The protein ratios of quantified and normalized proteins were processed in Microsoft Excel to calculate the group means and fold changes. All plasma samples were grouped into two sets; before and after exercise. Only proteins quantified in at least three horses from each haplotype group (EPH and SPH), i.e., proteins with at least three biological replicates in each haplotype group, were included in the analyses. Perseus version 1.6.15.20 was used to analyze differences between the groups, by applying t-test on log2 protein ratios. Five different comparisons were made; SPH and EPH at rest and during exercise as well as rest versus exercise within each haplotype. Additionally, paired ratio analysis was performed, to compare differences in protein levels during exercise vs. at rest for each haplotype i.e., (SPH- rest/SPH-exercise)/(EPH-rest/EPH-exercise). For each comparison, fold change values were calculated by dividing the average values for the compared groups and P-values ≤ 0.05 were considered to be significant. GO-annotations downloaded from the Uniprot database into Perseus were used to support the identification of changing biological processes. Furthermore, the GO enrichment analysis and visualization tool g:GOst from g:Profiler was used to identify the biological processes affected in each of the different contrasts (32).

## Declarations

### Ethics approval and consent to participate

Blood sample collection was approved by the ethics committee for animal experiments in Uppsala, Sweden (number: 5.8.18-15453/2017 and 5.8.18-01654/2020).

### Consent for publication

Not applicable.

### Availability of data and materials

The datasets used and analyzed during the current study will be deposited to the SRA.

### Authorś contributions

G.L. designed the project and led the genetic mapping together with J.R.S.M. M.K.R. was supervised with main input from G.L. and R.N. to perform the sampling, genetic mapping and blood pressure measurements. M.Å. designed genotyping assays. A.J. performed genotyping. M.K.R. and K.F. performed haplotype analysis. L.O. performed the mapDATAge analysis. R.N. designed the long-read sequencing experiment, performed ChIP- sequencing and proteomics data analysis. A.T. performed proteomic experiments and analysis. B.D.V. contributed with supervision of M.K.R. and A.R. contributed with mentoring of K.F. B.E .advised the blood pressure measurements. C.M.M. assisted with human comparisons and in the interpretation of the blood pressure results. G.A. performed transcription factor binding site analyses. J.R.S.M. assisted with the comparative genomic analysis and performed the transcription factor binding site analysis. K.F. lead the writing of the manuscript and created figures, with input from all co-authors.

## Acknowledgements

Thanks to all horse owners and trainers who participated in the study, The Swedish Trotting Association and the Norwegian Trotting Association for providing pedigree and performance data, Anna Darlene van der Heiden for the ONT sequencing, Tytti Vanhala for help with genotyping, Anna Johansson at the National Bioinformatics Infrastructure Sweden at SciLifeLab for bioinformatics advice, and the Department of Clinical Sciences Clinical training center (KTC) at SLU for providing the Cardell Veterinary monitor for the blood pressure measurements. Anna Svenssońs lab at the Department of Clinical Sciences SLU for running ELISA. Adnan Niazi at SLU Bioinformatics Infrastructure (SLUBI), Uppsala, Sweden SLUBI for supporting Nanopore data analysis. WGS was performed by the SNP&SEQ Technology Platform in Uppsala. The facility is part of the National Genomics Infrastructure (NGI) Sweden and Science for Life Laboratory. The SNP&SEQ Platform is also supported by the Swedish Research Council and the Knut and Alice Wallenberg Foundation. The computations and data handling were enabled by resources in project snic2018-8-226 provided by the Swedish National Infrastructure for Computing (SNIC) at UPPMAX, partially funded by the Swedish Research Council through grant agreement no. 2018-05973. The global relative quantitative proteomic analysis was performed at the Proteomics Core Facility, Sahlgrenska Academy at the University of Gothenburg, Gothenburg, Sweden.

