## Supplementary Table 2 for "An equine Endothelin 3 cis-regulatory variant links blood pressure modulation to elite racing performance"

| **Horse ID** | **Age (years)** | **Sex** | **Breed** | **Haplotype group** | **Exercise test** | **BP measurement** | **WGS** | **ONT seq** | **ELISA rest/exercise** | **Proteomics rest/exercise** |
| --- | --- | --- | --- | --- | --- | --- | --- | --- | --- | --- |
| 1 | 8 | S | CBT | SPH | + | R,E,P | NA | NA | +/+ | +/+ |
| 2 | 8 | S | CBT | SPH | + | R,E,P | NA | + | +/+ | +/+ |
| 3 | 4 | G | CBT | SPH | + | R,E,P | NA | NA | NA | NA |
| 4 | 5 | S | CBT | SPH | + | R,E,P | NA | + | +/+ | +/+ |
| 5 | 3 | G | CBT | SPH | + | R,E,P | NA | NA | +/+ | +/+ |
| 6 | 5 | M | CBT | SPH | NA | R | NA | NA | +/- | +/- |
| 7 | 5 | G | CBT | SPH | + | NA | NA | NA | +/+ | NA |
| 8 | 13 | G | CBT | SPH | + | NA | NA | + | +/+ | NA |
| 9 | 2 | S | CBT | SPH | + | NA | NA | NA | +/+ | +/+ |
| 10 | 11 | M | CBT | SPH | NA | NA | + | + | NA | NA |
| 11 | 5 | S | SB | SPH | NA | NA | + | NA | NA | NA |
| 12 | 8 | S | CBT | EPH | + | R,E,P | NA | + | +/+ | +/+ |
| 13 | 11 | S | CBT | EPH | + | R,E,P | NA | + | +/+ | +/+ |
| 14 | 6 | S | CBT | EPH | + | R,E,P | NA | NA | +/+ | NA |
| 15 | 9 | S | CBT | EPH | + | R,E,P | NA | NA | +/+ | +/+ |
| 16 | 7 | M | CBT | EPH | + | R,E,P | NA | + | +/+ | +/+ |
| 17 | 5 | M | CBT | EPH | + | R,E,P | NA | NA | +/+ | NA |
| 18 | 4 | G | CBT | EPH | + | R,E | NA | NA | +/+ | NA |
| 19 | 5 | G | CBT | EPH | + | R, P | NA | NA | +/+ | NA |
| 20 | 7 | M | CBT | EPH | + | R,P | NA | NA | +/+ | NA |
| 21 | 3 | S | CBT | EPH | + | R,P | NA | NA | +/+ | NA |
| 22 | 3 | S | CBT | EPH | + | R,P | NA | NA | +/+ | +/+ |
| 23 | 3 | S | CBT | EPH | + | R,P | NA | NA | +/+ | NA |
| 24 | 3 | S | CBT | EPH | + | R,P | NA | NA | +/+ | NA |
| 25 | 8 | G | CBT | EPH | + | R | NA | NA | +/+ | NA |
| 26 | 6 | S | CBT | EPH | NA | R | NA | NA | +/- | NA |
| 27 | 3 | S | CBT | EPH | NA | R | NA | NA | +/- | NA |
| 28 | 3 | M | CBT | EPH | NA | R | NA | NA | +/- | NA |
| 29 | 7 | S | CBT | EPH | NA | NA | NA | NA | +/- | NA |
| 30 | 2 | S | CBT | EPH | + | NA | NA | NA | +/+ | +/+ |
| 31 | 3 | M | CBT | EPH | NA | NA | NA | NA | +/- | NA |
| 32 | 3 | M | CBT | EPH | NA | NA | NA | NA | +/- | NA |
| 33 | 22 | S | CBT | EPH | NA | NA | + | + | NA | NA |
| 34 | 4 | S | SB | EPH | NA | NA | + | NA | NA | NA |
| 35 | 12 | S | CBT | HET | NA | R | NA | NA | NA | NA |
| 36 | 6 | S | CBT | HET | + | R, P | NA | NA | -/+ | NA |
| 37 | 10 | S | CBT | HET | + | P | NA | NA | +/+ | NA |
| 38 | 4 | S | CBT | HET | + | R | NA | NA | +/+ | NA |
| 39 | 3 | M | CBT | HET | + | R | NA | NA | +/+ | NA |
| 40 | 4 | G | CBT | HET | NA | R | NA | NA | +/- | NA |
| 41 | 5 | M | CBT | HET | NA | R | NA | NA | +/- | NA |
| 42 | 5 | M | CBT | HET | NA | R | NA | NA | +/- | NA |
| 43 | 4 | G | CBT | HET | + | R | NA | NA | +/+ | NA |
| 44 | 3 | G | CBT | HET | + | NA | NA | NA | -/+ | NA |
| 45 | 6 | S | CBT | HET | + | NA | NA | NA | -/+ | NA |
| 46 | 5 | M | CBT | HET | NA | NA | NA | NA | +/- | NA |

Supplementary Table 2. Information about all horses used in the various analyses.

S: Stallion, G: Gelding, M: Mare, CBT: Coldblooded trotter, SB: Standardbred,BP: blood pressure, R: at rest, E: during exercise, P: post-exercise, WGS: Whole-genome sequencing, ONT seq: Targeted MinION Oxford Nanopore sequencing, Proteomics: Global relative quantitative proteomic analysis, SPH: Horses homozygous for the sub-elite performing haplotype, EPH: Horses homozygous for the elite-performing haplotype, HET: Horses heterozygous.
