## Supplementary Table 4 for "An equine Endothelin 3 cis-regulatory variant links blood pressure modulation to elite racing performance"

|  |  |  |  | Binding Affinity | |  |  |
| --- | --- | --- | --- | --- | --- | --- | --- |
| SNP | BP | EVA 3 | SPH/EPH | SPH Allele | EPH Allele | Matrix_ID | Matrix name |
| 1 | 46713478 | rs395117226 | G/A | 0.0492 | 0.014 | MA0069.1 | Pax6 |
| 2 | 46715974 | rs397265747 | G/A | 0.0417 | 0.312 | MA0102.2 | CEBPA |
| 3 | 46717451 | rs396474304 | T/G | 0.00162 | 0.329 | MA0057.1 | MZF1_5-13 |
|  | 46717451 | rs396474304 | T/G | 0.0297 | 0.625 | MA0152.1 | NFATC2 |
|  | 46717451 | rs396474304 | T/G | 0.0455 | 0.645 | MA0056.1 | MZF1_1-4 |
|  | 46717451 | rs396474304 | T/G | 0.0109 | 0.0941 | MA0051.1 | IRF2 |
| 4 | 46717528 | rs69244081 | T/C | 0.0275 | 0.00295 | MA0017.1 | NR2F1 |
| 5 | 46717742 | rs69244084 | A/G | 0.00461 | 0.0639 | MA0038.1 | Gfi |
| 6 & 7* | 46717854/46717860 | rs69244085/rs69244086 | CC/TT | 0.744 | 0.0445 | MA0160.1 | NR4A2 |
|  | 46717854/46717860 | rs69244085/rs69244086 | CC/TT | 0.386 | 0.032 | MA0141.1 | Esrrb |
| 8 | 46717999 | rs69244088 | C/T | 0.00356 | 0.0255 | MA0133.1 | BRCA1 |
| 9 | 46718095 | rs69244089 | T/C | 0.035 | 0.00228 | MA0095.1 | YY1 |
|  | 46718095 | rs69244089 | T/C | 0.145 | 0.0376 | MA0057.1 | MZF1_5-13 |
| 10 | 46718361 | rs396281591 | C/T | NA | NA | NA | NA |
| 11 | 46718436 | rs394573286 | T/C | 0.0254 | 0.177 | MA0088.1 | znf143 |
| 12 | 46718895 | rs69244091 | G/A | 0.0389 | 0.283 | MA0105.1 | NFKB1 |
| 13 | 46718964 | rs69244093 | G/A | 0.912 | 0.0245 | MA0109.1 | Hltf |
| 13 | 46718964 | rs69244093 | G/A | 0.142 | 0.00755 | MA0083.1 | SRF |
|  | 46718964 | rs69244093 | G/A | 0.145 | 0.0103 | MA0158.1 | HOXA5 |
|  | 46718964 | rs69244093 | G/A | 0.136 | 0.0438 | MA0018.2 | CREB1 |
| 14 | 46719042 | rs69244095 | G/A | NA | NA | NA | NA |

Supplementary Table 3. Potential transcription factor binding changes with jaspar vertebrate matrices and human promoters.

* Both alleles implicated in matrix. Green indicates significant binding affinity.
