## Supplementary Table 5 for "An equine Endothelin 3 cis-regulatory variant links blood pressure modulation to elite racing performance"

Supplementary Table S4. Mean values ± SD for all blood pressure measurments.

| **Trait** | **EPH (n)** | **HET (n)** | **SPH (n)** | **P-value^1^** |
| --- | --- | --- | --- | --- |
| Age | 5.1 ± 2.2 (17) | 5.4 ± 2.9 (8) | 5.9 ± 2.8 (7) | 0.70 |
| Sex (%) |  |  |  |  |
| Mare | 23.5 (4) | 37.5 (3) | 14.2 (1) |  |
| Gelding | 29.4 (5) | 25.0 (2) | 42.9 (3) |  |
| Stallion | 47.0 (8) | 37.5 (3) | 42.9 (3) |  |
| Rest, before exercise |  |  |  |  |
| SBP | 123.1 ± 10.7 (17) | 116.9 ± 11.9 (8) | 121.4 ± 10.7 (7) | 0.43 |
| DBP | 75.1 ± 10.1 (17) | 75.1 ± 10.2 (8) | 77.9 ± 4.8 (7) | 0.78 |
| MAP | 93.2 ± 9.8 (17) | 92.0 ± 11.2 (8) | 95.3 ± 6.5 (7) | 0.79 |
| BPM | 35.8 ± 4.7 (17) | 37.4 ± 4.5 (8) | 38.2 ± 4.1 (7) | 0.46 |
| PP | 47.9 ± 7.6 (17) | 41.8 ± 6.1 (8) | 43.5 ± 9.7 (7) | 0.16 |
| During exercise (directly after uphill interval) |  |  |  |  |
| SBP | 156.5 ± 20.6 (7) | NA | 186.3 ± 17.2 (5) | **0.02** |
| DBP | 94.3 ± 19.1 (7) | NA | 119.9 ± 19.8 (5) | **0.05** |
| MAP | 116.0 ± 11.0 (7) | NA | 143.3 ± 21.6 (5) | **0.02** |
| BPM | 86.4 ± 13.6 (7) | NA | 97.5 ± 7.4 (5) | 0.13 |
| PP | 62.2 ± 28.7 (7) | NA | 66.3 ± 17.6 (5) | 0.78 |
| During exercise (five min after uphill interval) |  |  |  |  |
| SBP | 144.7 ± 12.3 (7) | NA | 189.3 ± 36.2 (5) | **0.01** |
| DBP | 92.3 ± 7.3 (7) | NA | 109.7 ± 24.6 (5) | 0.10 |
| MAP | 109.5 ± 5.3 (7) | NA | 140.2 ± 29.0 (5) | **0.02** |
| BPM | 77.0 ± 12.9 (7) | NA | 85.7 ± 14.2 (5) | 0.30 |
| PP | 52.4 ± 14.3 (7) | NA | 79.5 ± 23.2 (5) | **0.03** |
| Post-exercise  (10 min) |  |  |  |  |
| SBP | 132.9 ± 8.9 (9) | NA | 144.3 ± 3.4 (3) | 0.06 |
| DBP | 81.3 ± 11.4 (9) | NA | 91.3 ± 3.0 (3) | 0.18 |
| MAP | 101.9 ± 10.9 (9) | NA | 112.2 ± 4.7 (3) | 0.15 |
| BPM | 58.2 ± 10.6 (9) | NA | 57.8 ± 1.5 (3) | 0.95 |
| PP | 51.6 ± 5.9 (9) | NA | 53.0 ± 3.7 (3) | 0.71 |
| Post-exercise  (20 min) |  |  |  |  |
| SBP | 119.6 ± 9.6 (11) | NA | 133.0 ± 14.9 (5) | **0.05** |
| DBP | 71.5 ± 9.6 (11) | NA | 72.6 ± 11.8 (5) | 0.84 |
| MAP | 87.2 ± 9.3 (11) | NA | 96.7 ± 17.2 (5) | 0.17 |
| BPM | 51.0 ± 8.0 (11) | NA | 47.5 ± 5.9 (5) | 0.40 |
| PP | 48.2 ± 6.7 (11) | NA | 60.4 ± 7.5 (5) | **0.006** |
| Post-exercise  (30 min) |  |  |  |  |
| SBP | 119.8 ± 10.0 (8) | NA | 122.1 ± 14.1 (5) | 0.74 |
| DBP | 70.4 ± 11.6 (8) | NA | 68.9 ± 8.5 (5) | 0.82 |
| MAP | 88.4 ± 9.9 (8) | NA | 89.6 ± 11.0 (5) | 0.84 |
| BPM | 45.6 ± 7.7 (8) | NA | 45.0 ± 4.2 (5) | 0.87 |
| PP | 49.5 ± 6.7 (8) | NA | 53.2 ± 5.8 (5) | 0.33 |
| Post-exercise  (40 min) |  |  |  |  |
| SBP | 117.4 ± 13.3 (8) | NA | 115.8 ± 18.1 (4) | 0.86 |
| DBP | 68.0 ± 8.4 (8) | NA | 69.2 ± 16.5 (4) | 0.87 |
| MAP | 88.3 ± 9.6 (8) | NA | 88.0 ± 21.3 (4) | 0.96 |
| BPM | 42.6 ± 6.6 (8) | NA | 44.8 ± 2.6 (4) | 0.56 |
| PP | 49.4 ± 9.2 (8) | NA | 46.6 ± 3.1 (4) | 0.58 |

SBP: systolic blood pressure, DBP: diastolic blood pressure, MAP: mean arterial pressure, BPM: beats per min, PP: pulse pressure.

^1^ Tukey´s HSD test was performed
