## Supplementary Table 6 for "An equine Endothelin 3 cis-regulatory variant links blood pressure modulation to elite racing performance"

| **Sample ID** | **Haplotype** | **At rest (pg/ml)** | **CV (%)** | **During exercise (pg/ml)** | **CV (%)** | ***p*-value** |
| --- | --- | --- | --- | --- | --- | --- |
| 1 | SPH | 7.95 ± 1.61 | 20.3 | 7.43 ± 0.38 | 5.04 | 0.7 |
| 2 | SPH | 8.36 ± 0.06 | 0.74 | 10.47 ± 0 | 0.00 | **< 0.001** |
| 4 | SPH | 13.09 ± 0.09 | 0.65 | 12.42 ± 0.17 | 1.38 | 0.04 |
| 5 | SPH | 9.44 ± 0.98 | 10.36 | 7.78 ± 1.12 | 14.39 | 0.25 |
| 6 | SPH | 12.60 ± 0.60 | 4.74 | NA | NA | NA |
| 7 | SPH | 13.87 ± 0.34 | 2.44 | 13.26 ± 0.68 | 5.13 | 0.38 |
| 8 | SPH | 14.25 ± 2.00 | 14.04 | 13.90 ± 0.90 | 6.49 | 0.84 |
| 9 | SPH | 14.52 ± 0.60 | 4.07 | 14.94 ± 1.35 | 9.03 | 0.73 |
| 12 | EPH | 6.81 ± 0.38 | 5.55 | 6.50 ± 0.06 | 0.97 | 0.37 |
| 13 | EPH | 7.68 ± 0.09 | 1.16 | 7.18 ± 0.27 | 3.75 | 0.13 |
| 14 | EPH | 5.41 ± 0.11 | 2.01 | 5.64 ± 0.43 | 7.7 | 0.55 |
| 15 | EPH | 10.52 ± 0.06 | 0.58 | 6.45 ± 1.27 | 19.65 | **0.05** |
| 16 | EPH | 4.58 ± 0.98 | 21.45 | 6.05 ± 0.70 | 11.59 | 0.23 |
| 17 | EPH | 7.56 ± 0.31 | 4.94 | 6.41 ± 1.19 | 15.68 | 0.32 |
| 18 | EPH | 7.87 ± 1.24 | 15.8 | 5.6 ± 0.19 | 3.44 | 0.13 |
| 19 | EPH | 7.08 ± 0.11 | 1.5 | 8.94 ± 0.21 | 2.33 | 0.62 |
| 20 | EPH | 6.48 ± 0.75 | 11.57 | 8.20 ± 0.84 | 10.22 | 0.16 |
| 21 | EPH | 6.68 ± 2.57 | 26.11 | 8.41 ± 2.20 | 26.11 | 0.54 |
| 22 | EPH | 6.85 ± 0 | 0.00 | 9.01 ± 0.31 | 3.46 | **0.01** |
| 23 | EPH | 11.82 ± 0.69 | 5.81 | 10.54 ± 0.09 | 0.82 | 0.12 |
| 24 | EPH | 6.25 ± 0 | 0.00 | 7.01 ± 0 | 0.00 | NA |
| 25 | EPH | 7.56 ± 0.44 | 5.77 | 5.55 ± 1.03 | 18.54 | 0.13 |
| 26 | EPH | 8.49 ± 1.67 | 19.68 | NA | NA | NA |
| 27 | EPH | 9.81 ± 1.03 | 10.5 | NA | NA | NA |
| 28 | EPH | 13.39 ± 0.17 | 1.27 | NA | NA | NA |
| 29 | EPH | 10.39 ± 0.21 | 1.97 | NA | NA | NA |
| 30 | EPH | 10.84 ± 0.17 | 1.59 | 13.45 ± 0.09 | 0.63 | **0.003** |
| 31 | EPH | 5.48 ± 0.87 | 15.88 | NA | NA | NA |
| 32 | EPH | 7.00 ± 0.64 | 9.1 | NA | NA | NA |
| 36 | HET | NA | NA | 6.47 ± 0.96 | 0.00 | NA |
| 37 | HET | 7.08 ± 0.11 | 1.97 | 6.93 ± 0.11 | 1.53 | 0.29 |
| 38 | HET | 8.32 ± 2.93 | 35.21 | 5.25 ± 0.77 | 14.57 | 0.29 |
| 39 | HET | 9.81 ± 0.82 | 8.4 | 3.91 ± 0.00 | 0.00 | 0.01 |
| 40 | HET | 3.91 ± 0.00 | 0.00 | NA | NA | NA |
| 41 | HET | 3.91 ± 0.00 | 0.00 | NA | NA | NA |
| 42 | HET | 10.25 ± 1.23 | 12.02 | NA | NA | NA |
| 43 | HET | 6.40 ± 0.21 | 3.35 | 13.04 ± 2.52 | 19.3 | 0.07 |
| 44 | HET | NA | NA | 6.48 ± 0.11 | 1.65 | NA |
| 45 | HET | 6.17 ± 0.11 | 1.74 | NA | NA | NA |
| 46 | HET | 14.46 ± 2.10 | 14.51 | NA | NA | NA |

Supplementary Table 5. Plasma concentration of Endothelin 1 (EDN1) (mean + standard deviation) at rest and during exercise.

CV= coefficient of variation of the technical replicates

p-values generated from Student´s t-test, significant values in bold
