## Supplementary Table 7 for "An equine Endothelin 3 cis-regulatory variant links blood pressure modulation to elite racing performance"

| **Sample ID** | **Haplotype** | **At rest (pg/ml)** | **CV (%)** | **During exercise (pg/ml)** | **CV (%)** | ***p-*value** |
| --- | --- | --- | --- | --- | --- | --- |
| 1 | SPH | 20.51 ± 1.18 | 5.77 | 21.32 ± 0.41 | 1.93 | 0.38 |
| 2 | SPH | 21.15 ± 0.24 | 1.15 | 19.81 ± 0.52 | 2.61 | 0.7 |
| 4 | SPH | 17.59 ± 0.17 | 0.97 | 18.67 ± 0.76 | 4.05 | 0.19 |
| 5 | SPH | 19.03 ± 0.46 | 2.40 | 19.78 ± 1.6 | 8.00 | 0.52 |
| 6 | SPH | 8.99 ± 0 | 0.00 | NA | NA | NA |
| 7 | SPH | 19.97 ± 0.25 | 1.24 | 21.95 ± 0.68 | 3.12 | 0.06 |
| 8 | SPH | 17.39 ± 0.44 | 2.56 | 15.02 ± 0.28 | 1.86 | 0.41 |
| 9 | SPH | 13.74 ± 0.54 | 3.96 | 13.99 ± 0.63 | 4.52 | 0.7 |
| 12 | EPH | 22.98 ± 0.98 | 4.24 | 23.13 ± 1.83 | 7.91 | 0.85 |
| 13 | EPH | 23.76 ± 2.97 | 12.5 | 24.51 ± 2.16 | 8.80 | 0.87 |
| 14 | EPH | 24.61 ± 0.46 | 1.85 | 23.87 ± 0.33 | 1.37 | 0.2 |
| 15 | EPH | 24.12 ± 0.37 | 1.52 | 22.52 ± 0.22 | 0.97 | **0.04** |
| 16 | EPH | 30.08 ± 0.03 | 0.11 | 29.93 ± 4.43 | 6.21 | 0.12 |
| 17 | EPH | 23.54 ± 1.45 | 6.15 | 31.99 ± 0.67 | 2.10 | **0.02** |
| 18 | EPH | 22.46 ± 2.21 | 9.85 | 24.97 ± 0.97 | 3.89 | 0.47 |
| 19 | EPH | 21.09 ± 1.62 | 7.68 | 27.82 ± 4.74 | 17.03 | 0.18 |
| 20 | EPH | 21.48 ± 0.13 | 0.63 | 19.85 ± 0.68 | 3.45 | 0.08 |
| 21 | EPH | 26.38 ± 1.41 | 5.34 | 30.26 ± 0.31 | 1.02 | 0.06 |
| 22 | EPH | 20.12 ± 2.18 | 10.84 | 22.09 ± 0.47 | 2.12 | 0.33 |
| 23 | EPH | 17.20 ± 0.56 | 3.28 | 19.94 ± 1.50 | 7.54 | 0.13 |
| 24 | EPH | 11.45 ± 2.80 | 24.46 | 10.66 ± 1.38 | 12.91 | 0.76 |
| 25 | EPH | 23.87 ± 2.00 | 5.01 | 24.85 ± 1.03 | 4.15 | 0.07 |
| 26 | EPH | 28.17 ± 2.77 | 9.84 | NA | NA | NA |
| 27 | EPH | 16.34 ± 0.78 | 4.79 | NA | NA | NA |
| 28 | EPH | 22.81 ± 2.39 | 10.47 | NA | NA | NA |
| 29 | EPH | 15.99 ± 0.57 | 3.57 | NA | NA | NA |
| 30 | EPH | 23.67 ± 0.08 | 0.33 | 21.78 ± 0.44 | 2.04 | **0.03** |
| 31 | EPH | 19.79 ± 1.85 | 9.33 | NA | NA | NA |
| 32 | EPH | 23.73 ± 0.39 | 1.66 | NA | NA | NA |
| 36 | HET | NA | NA | 14.20 ± 0.66 | 4.63 | NA |
| 37 | HET | 18.23 ± 1.60 | 8.78 | 21.52 ± 1.28 | 5.93 | 0.09 |
| 38 | HET | 20.71 ± 0.54 | 2.62 | 21.88 ± 3.01 | 13.76 | 0.64 |
| 39 | HET | 22.37 ± 0.07 | 0.30 | 24.65 ± 1.43 | 5.81 | 0.15 |
| 40 | HET | 16.94 ± 1.48 | 8.76 | NA | NA | NA |
| 41 | HET | 22.22 ± 1.87 | 8.4 | NA | NA | NA |
| 42 | HET | 13.58 ± 0.22 | 1.63 | NA | NA | NA |
| 43 | HET | 20.66 ± 0.75 | 3.61 | 23.61 ± 3.16 | 13.36 | 0.33 |
| 44 | HET | NA | NA | 29.20 ± 1.90 | 4.06 | NA |
| 45 | HET | 20.09 ± 0.34 | 1.70 | NA | NA | NA |
| 46 | HET | 19.22 ± 0.34 | 1.79 | NA | NA | NA |

Supplementary Table 6. Plasma concentration of Endothelin 3 (EDN3) (mean + standard deviation) at rest and during exercise

CV= coefficient of variation of the technical replicates

p-values generated from Student´s t-test. significant values in bold
