## Supplementary Table 9 for "An equine Endothelin 3 cis-regulatory variant links blood pressure modulation to elite racing performance"

Supplementary Table 8. List of primers

| Forward primer | Sequence (5'-3') | Reverse primer | Sequence (5'-3') | Product size (bp) |
| --- | --- | --- | --- | --- |
| EDN3_1_F | CCAGCACCACCACTTCTTCT | EDN3_1_R | TCCCATGACTCCTTGTCCCT | 17774 |
| EDN3_2_F | TAGATTTCTTTACAAAGGGTCTGAAGT | EDN3_2_R | GGCCTAGAATCTTGCAACATCC | 10463 |
