## Supplementary Table 10 for "An equine Endothelin 3 cis-regulatory variant links blood pressure modulation to elite racing performance"

| **Breed** | **n** |
| --- | --- |
| Arabian horses | 30 |
| Thoroughbred | 30 |
| Standardbred | 45 |
| Coldblooded trotters | 224 |
| Finnhorses | 4 |
| Gotland pony | 3 |
| Shetland pony | 3 |
| Ardennes | 20 |
| Icelandic horses | 7 |
| North-Swedish draught | 18 |
| Exmoor pony | 23 |
| Fjord horse | 2 |
| Donkey | 4 |
| Przewalski | 3 |

**Supplementary Table 9. List of horses included in the MassArray genotyping analysis**
